# The transcription factor NO TRANSMITTING TRACT / WIP2 regulates cytokinin homeostasis in Arabidopsis

**DOI:** 10.1101/2025.04.03.645522

**Authors:** David Díaz-Ramírez, Edgar Demesa-Arevalo, Yolanda Durán-Medina, Rosa E. Becerra-García, José E. Cruz-Valderrama, Humberto Herrera-Ubaldo, Ricardo A. Chávez Montes, Maurizio Di Marzo, Lucia Colombo, Martin Sagasser, Rüdiger Simon, Ondrej Novak, Stefan de Folter, Nayelli Marsch-Martínez

## Abstract

NO TRANSMITTING TRACT / WIP2 (NTT/WIP2) is a zinc finger transcription factor that alters developmental programs in Arabidopsis when overexpressed. Loss of *NTT* function, alone or in combination with closely related genes, leads to developmental defects in gynoecia and roots, respectively. Its overexpression phenotype resembles the effects of cytokinin treatments, suggesting a potential connection to this hormonal pathway. To further investigate this connection, we conducted cytokinin content and response evaluations, and transcriptome analyses in an inducible NTT line. Alterations in cytokinin content and response, as evaluated with the *TCSn::GFP* line, were observed. In both cases, as in the global expression analyses, the observed changes were dependent on the tissue analyzed. Moreover, Y1H and transactivation assays suggested direct NTT binding to the *IPT5*, *AHP6* and *CKX7*, regulatory regions. We propose that NTT, among other mechanisms, regulates development through the modulation of cytokinin homeostasis at different levels, including its local concentration. Since *NTT* is a target of AUXIN RESPONSE FACTOR 5 / MONOPTEROS, the results presented here indicate that NTT connects an auxin input to a change in cytokinin levels.

## Introduction

The transcription factor WIP2 / NO TRANSMITTING TRACT (NTT) belongs to the WIP subfamily of zinc finger proteins that can localize in the nucleus and share a domain comprising four zinc fingers in tandem, which initiates with the amino acids Trp, Ile, Pro (WIP, hence the name, Sagasser *et al*., 2001). This subfamily includes six members in *Arabidopsis thaliana* (Appelhagen *et al.,* 2010). *WIP2/NTT* is expressed in different tissues during plant development, including the early embryo, shoot apical meristem (SAM), root apical meristem (RAM), vasculature and gynoecium, among others (Crawford *et al*., 2007, 2015; Chung *et al*., 2013; Herrera-Ubaldo et al., 2019; Marsch-Martínez *et al*., 2014).

Loss and gain of function alleles of *WIP2/NTT* in Arabidopsis have revealed several functions. During the development of the gynoecium, the female reproductive organ in flowers, WIP2/NTT participates in the establishment of the transmitting tract, an internal tissue that facilitates pollen tube growth through the ovary to reach and fertilize ovules. Loss of *WIP2/NTT* function affects the development of this tissue causing partial sterility. It was therefore named *NO TRANSMITTING TRACT (*Crawford *et al.,* 2007) and will be addressed as *NTT* through the rest of the text for simplicity. Alterations in *NTT* expression also affect replum and valve margin development (Chung *et al*., 2013; Marsch-Martínez *et al*., 2014). NTT, in combination with two other WIP members, WIP4 and WIP5, are key regulators of embryo development and RAM establishment. In the absence of these three proteins, seedlings lack roots (Crawford *et al., 2015*).

Although mutant phenotypes indicate that NTT regulates developmental pathways, we are just starting to elucidate its mechanisms of action. Most NTT targets have been studied in the context of gynoecium and fruit development (Marsch-Martinez *et al*., 2014; Herrera-Ubaldo *et al*., 2019). NTT is able to physically interact with many key gynoecium and fruit development regulators, such as REPLUMLESS (RPL), FRUITFUL (FUL), SHATTERPROOF1 and 2 (SHP1 and 2), SEEDSTICK (STK) and others (Marsch-Martinez *et al*., 2014; Herrera-Ubaldo *et al.,* 2019). Together, STK and NTT regulate genes involved in cell wall polysaccharide metabolism and lipid synthesis or transport, and some of them lead to altered gynoecium phenotypes when mutated (Herrera-Ubaldo *et al*., 2019).

Ren *et al*. (2018) explored the function of a *WIP* gene in *Gerbera hybrida*, finding that it acts as a transcriptional repressor and represses cell expansion in petals (Ren *et al*., 2018). In Arabidopsis, ChIP-seq and RNA-seq approaches using inducible lines of *NTT* and related gene *WIP1/TRANSPARENT TESTA 1 (TT1)*, gave more insights into binding sites and targets of these two WIP transcription factors (Gomez-Roldan *et al.,* 2020).

The phenotype of *ntt wip4 wip5* (*nww*) triple mutant seedlings is similar to the *monopteros* (*mp*) mutant phenotype, and the expression of *NTT* is lost in *mp.* MP is AUXIN RESPONSE FACTOR 5 (ARF5), a transcription factor that mediates the transcriptional response to auxin (Crawford *et al*., 2015). ChIP assays revealed that MP binds to auxin response elements (AuxREs) in the *NTT* genomic region (and similar regions in WIP4 and WIP5), suggesting that these WIPs act downstream the auxin response. They are not required for the initial activation of the DR5 auxin response marker during embryo development. However, DR5 expression is reduced at later stages, and a pulse of auxin can rescue the rootless phenotype in *nww* mutants, although these roots develop in a highly abnormal way (Crawford *et al*., 2015).

On the other hand, NTT physically interacts with cytokinin type-A Arabidopsis Response Regulators (ARR4 and ARR16; Dortay *et al*., 2008). Moreover, NTT also interacts with SHOOTMERISTEMLESS (STM) and binds to the promoter of *BREVIPEDICELLUS* (*BP*) (Marsch-Martinez *et al*., 2014), and both transcription factors have been reported to activate the cytokinin pathway (Yanai *et al*., 2005). In addition, cytokinin treatments rescue replum development in otherwise replum-less *ntt replumless (rpl)* double mutants (Zuñiga-Mayo *et al*., 2018), suggesting that cytokinins may be related to or play a role in NTT function.

Cytokinins are phytohormones involved in the regulation of many developmental processes, such as cell proliferation, root differentiation, leaf expansion, and senescence, among others (Hwang *et al*., 2012; Marquez-Lopez *et al.,* 2019; Wybouw and De Rybel, 2019). Cytokinins are synthesized mainly by ISOPENTENYL TRANSFERASES (IPT) and LONELY GUY (LOG) enzymes (Kuroha *et al.,* 2009; Miyawaki *et al*., 2006; Schaller *et al*., 2015; Duran-Medina *et al*., 2017; Wybouw and De Rybel, 2019). Cytokinins are then sensed by histidine kinases (AHKs), which autophosphorylate and then transfer the phosphate to histidine phosphotransferases (AHPs) (Hwang *et al.,* 2012; Kieber and Schaller, 2018). The AHPs then transfer the phosphate to the type-B ARRs, the response regulators that promote the transcription of the genes in response to cytokinin, including the negative regulators type-A ARRs (Kieber and Schaller, 2018). AHP6 is an exception because it does not transfer the phosphate and acts as a negative regulator of the pathway (Mähönen *et al*., 2006). Cytokinin inactivation is carried out through degradation by cytokinin oxidases (CKXs) which catalyze an irreversible oxidative breakdown (Werner *et al*., 2001, 2003), and by N- or O-glucosylation by cytokinin glucosyl transferases (Chen *et al.,* 2021). Degradation and N-glucosylation are irreversible, while O-glucosylation is a reversible inactivation process (Chen *et al.,* 2021).

Given the key role that cytokinins play in development, the central question of this work was to find out whether NTT regulates cytokinin genes. We found that NTT regulates genes that participate at different steps of the cytokinin pathway, influencing cytokinin content and response.

## Materials and methods

### Plant material, growth conditions and genotyping

The *Arabidopsis thaliana* lines used in this study are all in the Columbia (Col-0) background. Plants were grown under greenhouse or growth chamber conditions (22 °C, 16 hours of light) in soil or *in vitro,* in ½ MS medium supplemented with 0.5 % sugar and 1 % agar. The following lines were used in this work: the dexamethasone inducible line *35S::NTT-GR*, the recombineering line *gNTT-YPET, nww* triple mutant (Crawford *et al*., 2015), *35S::NTT (*Marsch-Martínez *et al*., 2014)*, pIPT5::GUS* (Dello Ioio *et al.,* 2007)*, pAHP6::GFP* (Mähönen *et al.,* 2006), *ckx7* (GABI-Kat; Kleinboelting *et al.,* 2012), *TCSn::GFP* (Zürcher *et al.,* 2013) and we generated *35S::NTT-GR ckx7, 35S::NTT-GR x pIPT5::GUS, 35S::NTT-GR x pAHP6::GFP* and *35S::NTT-GR x TCSn::GFP* through genetic crosses. Primer sequences for genotyping are in Table S1. The *Nicotiana benthamiana* plants used for the NanoLuc transactivation assays were grown in a 16 h light/8 h dark photoperiod at 25 °C for six weeks.

### Induction of NTT activity and cytokinin treatments

Most NTT inductions were performed on *in vitro*-grown *35S::NTT-GR* seedlings, at four days after germination (dag) by directly applying the solutions to the plants on the plates and removing the liquid after two minutes. 100 µM dexamethasone (DEX), 100 µM DEX + 30 µM cycloheximide (CHX), or mock solutions were used (Fig. S2). The DEX solution was prepared from a sterile 20 mM stock, first dissolved in ethanol, and then diluted with sterile water. To evaluate the induction response in the crosses, seeds were germinated directly in solid medium supplemented with 10 µM DEX. For induction under cytokinin treatments, seedlings were transplanted to medium containing 10 µM DEX and 10 µM 6-benzylaminopurine (BAP, dissolved in NaOH) or mock medium. For cytokinin treatments, four dag Col-0 seedlings (shown in Fig. 1) were transplanted onto medium containing 10 µM BAP.

**Figure 1.**
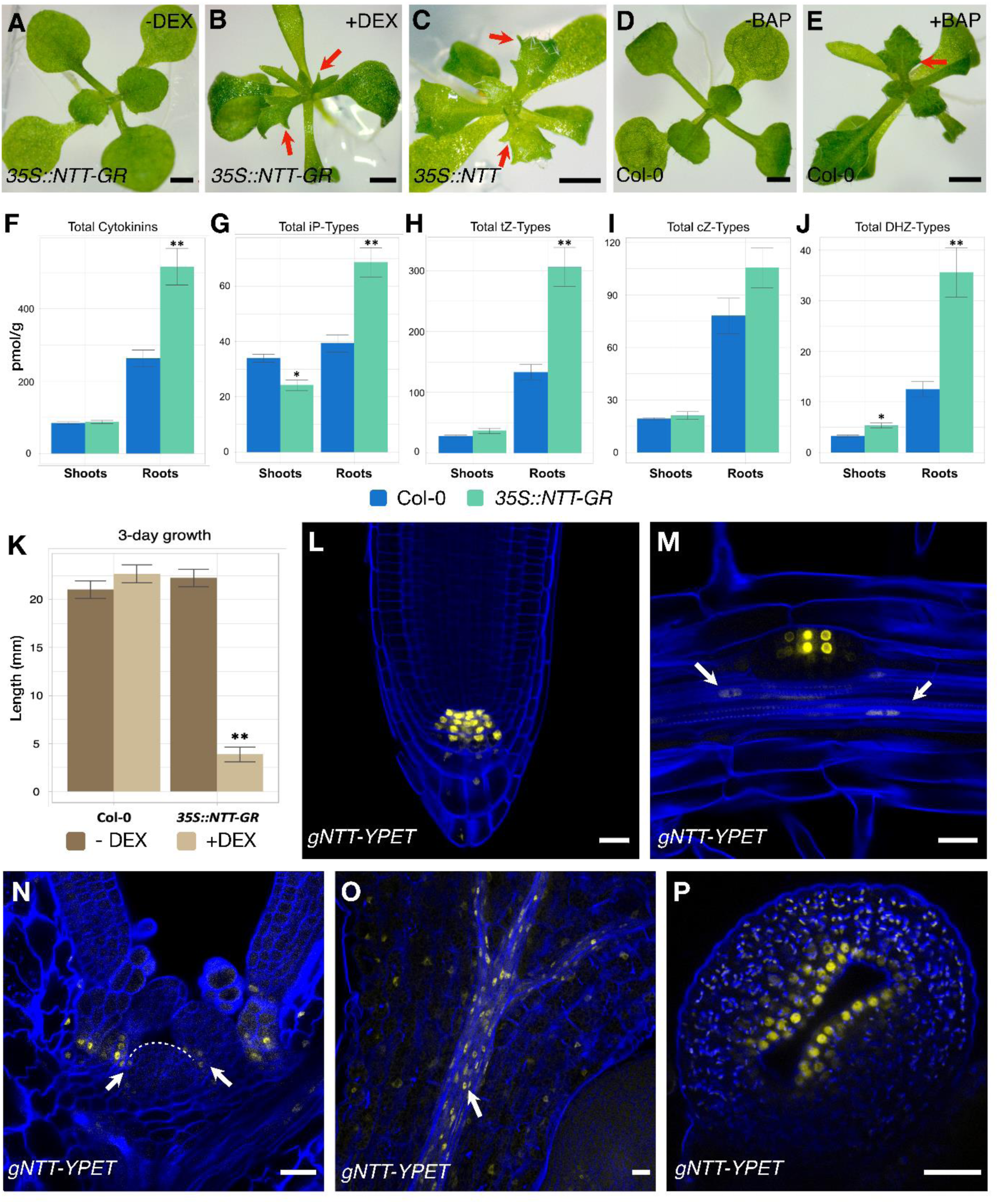
*NTT* OE phenotypes, cytokinin quantification and protein localization. (A-E) Representative images of 11 dag seedlings: (A) Mock treated *35S::NTT-GR* line. (B) DEX induced *35S::NTT-GR.* (C) NTT overexpression line (*35S::NTT* line). (D) Mock treated Col-0 seedling. (E) Col-0 seedlings after BAP treatment. Red arrows indicate leaf margins, and the irregularities observed under treatments or NTT overexpression. (F-J) Quantification of different types of cytokinin content in 6 dag Col-0 and *35S::NTT-GR* seedlings (root and shoot tissues were collected and analyzed separately, 2 dai, 4 dag seedlings): (F) Total cytokinins (ck), (G-J) Total (ribosides, glucosides, and free bases) (G) isopentenyl adenosine (iP) type ck; (H) trans-zeatin (tZ) types; (I) cis-zeatin (cZ) types; and (J) Dihydro-zeatin (DHZ) types. (K) Differences in root growth between mock and Dex treated Col-0 and *35S::NTT-GR* seedlings. Only the DEX treated *35S::NTT-GR* line is different from the rest. (F-K) Student’s t-test, * p ≤ 0.01, ** p ≤ 0. 001. (L-P) *gNTT-YPET* accumulation pattern: (L) in quiescent center, initials and columella in RAM; (M) In initiation of lateral roots and companion cells (white arrows) in root vasculature; (N) surrounding SAM (boundary cells, white arrows) and primordia (1-2 dag seedlings); (O) in companion cells in aerial vasculature (white arrow) and leaf epidermis; and (P) in the carpel marginal meristem (CMM), in a stage 6 gynoecium. L-O: 5 dag seedlings. Counterstaining L-O: SR2200, P: PI. Scale bars 1mm (A-E) and 20 µm (L-P).

### Cytokinin quantification

Quantification of CK metabolites was performed as described in (Svačinová *et al*., 2012). Four dag seedlings (Col-0 or *35S::NTT-GR*) were treated with DEX or mock as described previously. After a week of treatment, around 30 mg of shoots and roots of the treated and mock seedlings were harvested separately (four independent replicates), flash-frozen in liquid nitrogen, and stored at -80°C. Extraction, purification and quantification were done as described in (Duran-Medina *et al*., 2025; Svačinová *et al.,* 2012).

### RNA extraction

Aerial tissue (shoots) and roots of the treated seedlings were harvested separately for RNA extraction 30 min after the treatment with DEX and CHX, or eight hours after the treatment with DEX, and their respective controls were also harvested at the same times (three replicates, around 10 seedlings for aerial tissue, and around 20 for root tissue). RNA was extracted and purified using *Quick RNA* miniprep (Zymo Research, Orange, CA, USA). RNA quality assessment, and library construction were performed by Macrogen (Seoul, South Korea).

### RNA-seq, alignment, quantification, and differential expression analysis

RNA-seq was carried out at Macrogen using the TruSeq Stranded mRNA LT Sample Prep Kit and Illumina 100 base paired-end sequencing. The data was deposited at the European Nucleotide Archive under the study accession number PRJEB40086. After quality control using FastQC (https://www.bioinformatics.babraham.ac.uk/projects/fastqc/), pseudo-alignment and quantification in paired-end strand-specific mode versus the Arabidopsis Araport11 (Cheng *et al*., 2017) cDNA sequences (version 2016) was done using Kallisto (version 0.46.1; Bray *et al*., 2016). Kallisto data gene-level summarization and pre-processing was done with txt import (Soneson *et al*., 2016), and differential expression analysis was done with edgeR (Robinson *et al*., 2009; McCarthy *et al*., 2012) under R (R Core Team, 2022; https://www.R-project.org/).

### Gene Ontology categories enrichment analysis

Gene Ontology biological process categories enrichment analysis was done using the R package SetRank (“SetRank: Advanced Gene Set Enrichment Analysis. R package version 1.1.0. https://CRAN.R-/package=SetRank”; Simillion *et al*., 2017) using the Gene Ontology gene association file for Arabidopsis (dated 01-02-2019, available at the Gene Ontology Consortium website (Ashburner *et al.,* 2000; The Gene Ontology Consortium, 2019) in SetRank format as the annotation file, the complete p-value-ranked edgeR tables as the query, and all expressed genes in either root or aerial tissue as the universe. We considered a gene as expressed if it had a value of at least 1 TPM across all samples of the corresponding tissue.

### RT-qPCR

Confirmation of gene expression was made by RT-qPCR, using the One-Step RT-qPCR Kit (Nzytech) in a Step One^TM^ thermocycler (Applied Biosystems), including 3 biological and technical replicates (tissue was collected in the same conditions as for the RNAseq experiment). Expression levels were normalized to *ACTIN2.* Relative expression was calculated with the ΔΔCt method (Livak and Schmittgen, 2001). The primers used are listed in Table S1.

### GUS Staining

Samples were collected and stained in GUS solution (0.5% triton, 2.5 mM potassium ferricyanide, 2.5 mM potassium ferrocyanide, 1 M EDTA and 2 mM X-Gluc in 50 mM NaPO4), at 37 °C overnight. Afterwards, plants were cleared in 70 % ethanol to easily distinguish coloration.

### Whole-mount immunostaining

Immunostaining was performed as follows: 1 dag seedlings were fixed as described by Tran *et al*. (2021) but using MTSB instead of PBS (Herrera-Ubaldo *et al*., 2023). Cuticle solubilization was carried out following Pasternak *et al*. (2015). Samples then were digested for 27 minutes at 22°C, as in Tran *et al*. (2021), followed by permeabilization according to Pasternak *et al*. (2015). After this, samples were blocked for 30 minutes in 4% BSA before incubation with the primary antibodies (ab) overnight. The protocol was continued as described by Tran *et al*. (2021) for days 2 and 3 (washing and incubation with secondary ab). Anti-trans-zeatin riboside (Cat. 004 0312, Olchemim) or anti-N6-isopentenyladenosine (Cat. 004 0172, Olchemim), both raised in rabbit, were used as primary ab. As a technical control, we used Pol II S2p primary ab, raised in mouse (Cat. C15200005, Diagenode; Fig. S7E). Secondary ab used were Alexa Fluor plus 488 (anti-mouse, Cat. A32723, Invitrogen) and Alexa Fluor 568 (anti-rabbit, Cat. A-11036, Invitrogen).

### Imaging

Confocal imaging was obtained using a Zeiss LSM800 or LSM880 confocal microscope. For *gNTT-YPET*, seedlings were cleared using the ClearSee protocol (Kurihara *et al*., 2015) and stained with SCRI Renaissance 2200 (SR2200). Renaissance imaging was performed using a 405 nm laser for excitation (ex), with emission(em) detected between 410–511 nm. For YPET fluorescence ex: 514 nm, em: 519–541 nm. For *35S::NTT-GR x TCSn::GFP* and *35S::NTT-GR x pAHP6::GFP* crosses, seedlings were directly mounted in propidium iodide (PI, 1 µg/ml). PI ex: 561 nm, em: 617–700 nm. GFP (ex: 488 nm, em 525– 570 nm). For immunostaining, Alexa Fluor 568 ex: 561 nm, em: 586–626 nm. Alexa Fluor 488 ex: 488 nm em: 506–537 nm. All confocal imaging was performed using multi-tracking to detect each fluorophore independently. For phenotyping and GUS expression pattern analyses, images were acquired with a Zeiss Stemi 2000-C stereo microscope, coupled with axiocam ERc 5s camera (Carl Zeiss).

### Y1H assays

Binding of NTT to the promoter regions of putative target genes was tested in a Yeast one-hybrid system (Y1H), based on the Matchmaker Gold Yeast One-Hybrid protocol (https://www.takarabio.com). Promoter regions of *IPT5, KAN1, CKX7, AHP6* and *PUP10* (Fig. 7A) were amplified using primers indicated in Table S1, cloned in an Entry vector (pENTR-D topo, Invitrogen), and recombined with CZN1810, a Gateway-compatible version of pAbAi (Clontech; Danisman *et al*, 2012)., the next steps were done as described in Duran-Medina *et al*., 2025. The background growth for each bait clone is shown in Fig. S5.

NTT as prey clone (AD-NTT, pDEST22, Invitrogen; Lozano-Sotomayor *et al*., 2016) in the haploid PJ69-A yeast was mated with the bait clones and grown in YPAD medium. Diploid yeasts (promoter + AD-NTT) were selected on SD GLUC medium lacking tryptophan and uracil.

To test protein-DNA interactions, diploid yeasts representing the possible combinations were plated on SD GLUC medium lacking tryptophan and uracil, with different concentrations of AbA, 250 ng mL^-1^ for *KAN1, CKX7, IPT5-A, PUP10* and *IPT5*; 300 ng mL^-1^ for *AHP6-A*; 700 ng mL^-1^ for *AHP6*. Yeast growth was scored on the sixth day of incubation at 30°C.

### Assays of NanoLuc activity in plant extracts

The same promoters of *IPT5, CKX7,* and *AHP6* used for Y1H were cloned into pGWB601-NL3F10H (Urquiza-García and Millar, 2019) using LR clonase mix (Invitrogen), to generate the reporter constructs *pAHP6::NL, pIPT5::NL,* and *pCKX7::NL*, which then were transformed into *Agrobacterium tumefaciens* GV3101. The assays were conducted following the methodology described by Becerra-García *et al*. (2023).

### Statistical Analyses

For all comparisons made, like root length, cytokinin contents, RT-qPCR, and NanoLuc, statistical analysis were performed using the Student’s t test (*p < 0.05, **p < 0.01, ***p < 0.001). All plots were made using R Studio (R Studio Team, 2020; http://www.rstudio.com/).

## Results

### NTT alters plant shape and is expressed in roots and shoots

NTT constitutive overexpression alters the architecture of Arabidopsis organs and plants. One of these alterations is the development of irregular leaf margins, such as the formation of pronounced serrations or projections at the leaf borders (Fig. 1C). Similarly, induction of NTT activity by dexamethasone (DEX) in a *35S::NTT*-GR line, leads to increased serrations or projections in leaf margins that are visible one week after induction (Fig. 1A, B). Besides the leaf phenotypes, a more immediate evident effect is the growth inhibition of the main root (Fig. 1K) and increased lateral root development (Fig. S1). The induced phenotype lasts for days but is not permanent. Many plants recovered a wild-type phenotype two weeks after the induction. Then, only about 40% of the induced plants still showed severe phenotype alterations. We considered that the induction phenotypes presented some similarities to cytokinin treated Arabidopsis seedlings (Fig. 1D, E). This contributed to the question of whether the cytokinin pathway was among the processes or pathways that were affected and possibly regulated by NTT.

### NTT alters CK content in seedlings

To obtain information about cytokinin levels in the NTT overexpressors, we compared the cytokinin content between DEX-treated wild-type and DEX-induced NTT-GR plants. We used plants that had been induced for 1 week and analyzed shoots and roots separately. We found clear changes in cytokinin content after NTT induction, and interestingly, the changes in CK content in roots and shoots were different (Fig. 1 F-J, Table S2). Total cytokinin content increased evidently in induced roots, and the total cytokinin species that increased were mostly ribosides, nucleotides, and glucosides. In shoots, fewer statistical differences in total cytokinin content were detected. When different cytokinin types were analyzed (table S2), again, most changes were observed in the root tissue. There, total isopentenyl (iP), trans-zeatin (tZ) and dihydrozeatin (DHZ) cytokinins were increased when the induced NTT-GR was compared to DEX-treated wild-type roots. From them, the content of almost all iP, tZ and DHZ derivatives (ribosides, nucleotides, and glucosides) increased, as also occurred with the biologically active tZ and DHZ. The exceptions were free iP, iP9G, and DHZ9G, which did not present a statistically significant change. From all the cytokinins analyzed, cis-zeatin (cZ) and all its quantified derivatives remained unchanged in the root. In the shoot, cZ riboside some DHZ glucosides content increased, while iP glucosides (iP7G and iP9G) diminished. In summary, NTT induction caused different effects in cytokinin content in the distinct tissues, mainly an increase on cytokinin levels in root tissues.

### NTT induction causes general and specific changes in gene expression

To search for genes related to the cytokinin pathway, and to obtain a global view of the genes that could be regulated directly or indirectly by NTT, we performed comparative RNA-seq analyses. In young plants, *gNTT-YPET* reporter lines (Crawford *et al*., 2015) revealed *NTT* expression at the root tip (Fig. 1L, incipient and very young lateral roots (Fig. 1M), and in the regions around the shoot apical meristem (Fig. 1N) and in vasculature of the hypocotyl, cotyledons (Fig. 1O), roots (Fig 1M, arrows) and leaves (Chung *et al.,* 2013; Marsch-Martinez *et al*., 2014; Crawford *et al*., 2015; Herrera-Ubaldo *et al*., 2019). Expression at the carpel marginal meristem was also observed (Fig. 1P).

However, despite expression in these tissues, due to redundancy (Crawford *et al*., 2015; Du et al., 2022), the loss of *NTT/WIP2* function visibly affects only specific tissues of the gynoecium (transmitting tract and replum, Crawford et al., 2007, Marsch-Martínez *et al*., 2014), and leaf size at specific developmental stages (Diaz-Ramirez et al., 2022). Moreover, the triple *nww* mutant is severely affected, since very young plants lack roots and can only be recovered by auxin treatments (Crawford *et al*., 2015).

To avoid redundancy issues, an *NTT-GR* inducible line was used for the RNA-seq analyses. Moreover, since NTT is present in different tissues, we analyzed aerial and root tissues separately to obtain information about the regulation it exerts in these different contexts. We were also interested in investigating the regulation over time, discriminating between “early response genes”, putatively direct targets, and “late” or secondary targets.

The experiment was carried out using *35S::NTT-GR* seedlings four days after germination (dag), treated with a DEX or with a mock solution. Roots and shoots were collected 30 min or 8 h after DEX treatment, according to the table in Fig. S2. To reduce the chance of downstream regulation of genes by transcription factors that were induced or repressed by NTT, the translation inhibitor cycloheximide was included in the treatment of the samples that were collected at 30 min. Differential gene expression analyses were done between induced and the non-induced samples for each tissue and each time point (Fig 2A). 30 min after NTT induction we found 1546 differentially expressed genes in aerial tissue and 1302 in roots, and at 8 h we found 1057 and 1945, respectively (Fig 2A). Some DEGs were common in different tissue types and times, and 49 genes were found to be differentially expressed in all conditions.

**Figure 2.**
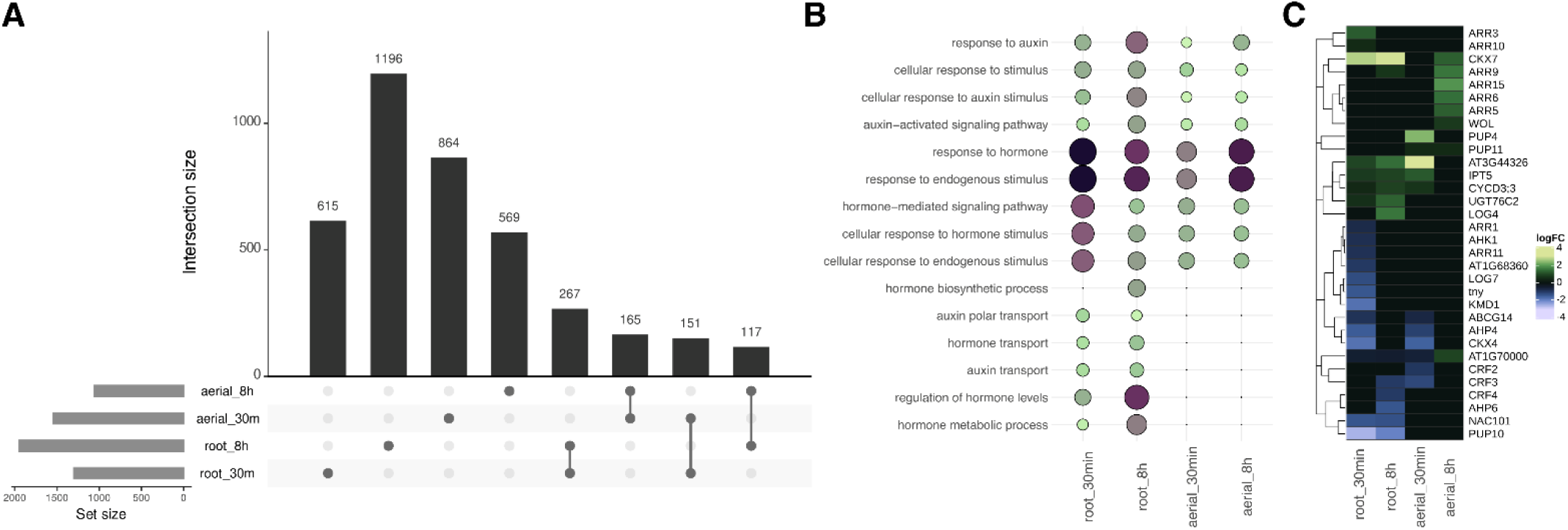
Analyses of differentially expressed genes (DEGs) obtained through RNA-seq analysis of roots and roots 30 minutes and 8 hours after NTT induction. (A) UpSet plot of DEGs found in the different conditions. Set size indicates the number of DEGs in shoots (aerial) and roots at 30 min or 8 hours after induction. Intersection size indicates the number of DEGs in the corresponding intersection. Only DEGs exclusive of a comparison or DEGs shared between same tissues or same induction times comparisons are shown. (B) Bubble plot of hormone-related GO Biological Process categories enrichment analysis. Bubble size and color represent the false discovery rate, with larger and darker points indicating lower FDR values. (C) Heatmap of the log2FC values for cytokinin-related DEGs. The full DEGs dataset is presented in Supplementary Dataset S1.

We noticed that in the case of roots there was a greater number of downregulated than upregulated genes. Conversely, in the aerial tissue there were more upregulated genes, about 70% of the genes, while almost 60% of the genes were downregulated in roots 30 min after induction (Dataset S1). Considering only DEGs with a log2 fold change of 2 and greater, these percentages increase, making more than 90% upregulated in aerial tissue (8 h), contrasting with more than 80% downregulated in roots (at 30 min).

Some of the genes that were found to be differentially expressed are genes related to development, such as *CUP SHAPED COTYLEDON 1* and *2* (*CUC1* and *2,* Aida *et al.,* 1997), the knotted1-like genes *KNAT1*, *4*, *6* and *7 (*Hake *et al*., 2004; Bharathan *et al*., 1999), *ASYMMETRIC LEAVES 2 (AS2,* Semiarti *et al*., 2001), *KANADI (KAN,* Bowman *et al.,* 2001*), WUSCHEL-related homeobox 4 (WOX4,* Ji *et al*., 2010*), SHORT ROOT (SHR,* Helariutta *et al*., 2000), and *NGATHA3 (NGA3,* Trigueros *et al.,* 2009), among others, and are very interesting as candidate NTT targets (Fig. S3B). Most were up or downregulated in specific tissues or at specific times, except for SHR, downregulated in 3 comparisons, and KAN, downregulated in all 4 comparisons. Interestingly, NTT itself was also found to be downregulated after induction in three comparisons (Fig S3B).

Next, a gene ontology (GO) enrichment analysis was done using both the complete lists and separating the up and downregulated genes. When exploring enrichment of DEGs belonging to certain GO categories in the differentially expressed gene datasets, we noticed that some were only enriched in the root or aerial tissue datasets, or at those obtained at earlier or later times, and others were found to be enriched in all (Dataset S2). Some of these categories included genes related to different hormonal pathways, from ethylene to auxin, gibberellin, jasmonate, abscisic acid, salicylic acid and cytokinin (Fig. 2B, S3A).

### NTT affects the expression of cytokinin genes

We then explored the possible regulatory role of NTT in the cytokinin pathway. Moreover, considering that *NTT* is a target of the ARF5/MP, and that the auxin and cytokinin pathways are frequently related, we also searched for auxin-related genes (Fig. S3C), besides cytokinin-related genes (Fig. 2C) in the DEG lists of roots and shoots at the two times. The DEG lists contained several genes corresponding to these hormonal pathways, and were related to biosynthetic, transport, signaling and response activities. For most of them, NTT affected their expression in a tissue and time-after-induction manner. The auxin-related genes included those responsible for the transcriptional response to auxin, Auxin Response Factors, and their interactors Aux/IAA proteins (Roosjen *et al*., 2018; Powers and Strader, 2019). Interestingly, *ARF5/MP* was found to be upregulated in the root 8 h after NTT induction. Moreover, besides *Aux/IAA* genes, differences in expression were detected in many genes belonging to the two other classes of early/primary auxin response genes, *Gretchen Hagen 3 (GH3)* and *Small Auxin Upregulated RNAs* (*SAURs)* (Hagen and Guilfoyle, 2002). Finally, genes coding for the auxin receptors AUXIN-SIGNALING F-BOX (AFB) 2 and 3 (Prigge *et al*., 2020), for the PIN1, 3, 5 and PILs transporters, and for the biosynthetic enzyme YUCCA9, also changed their expression in some of the tissues after NTT induction. Most changes in auxin gene expression were from low to moderate, with *IAA5, GH3.1* and *GH3.3* being the most upregulated of the auxin-related genes, while the most downregulated genes were *SAURs*.

On the other hand, some cytokinin-related genes presented a greater change in expression upon NTT induction (Fig. 2C). For this pathway, again genes involved in different processes were differentially expressed, some were up and some downregulated in a tissue and time-specific fashion, indicating that the effect of NTT in this pathway is complex. These genes included the *WOL/AHK4* cytokinin receptor (Mähönen *et al*., 2000; Yamada *et al.,* 2001), the downstream phosphotransfer or pseudo-phosphotransfer proteins *AHP4* and *AHP6* (Mähönen *et al*., 2006), and the transcriptional regulators type-B *ARRs ARR1, ARR10* and *ARR11* (Hill *et al*., 2013). The expression of the negative regulator *KISS ME DEADLY 1 (KMD1*), that targets type-B ARRs for degradation (Kim *et al*., 2013), and some type-A ARR negative regulators (To *et al*., 2004), including ARR5, 6 and 15, whose repression is important for SAM development (Liebfried *et al*., 2005), was also altered. Cytokinin transporter genes such as ABCG14 (Zhang *et al*., 2014) and PUPs (reviewed in Durán-Medina *et al*., 2017) were also affected. Cytokinin metabolism genes also presented changes in expression. Related to biosynthesis, we found *ISOPENTENYL TRANSFERASE 5* (*IPT5*; Miyawaki *et al.,* 2004) and *LONELY GUY 4* and *7* (*LOG4* and 7, Tokunaga *et al*., 2012). Related to cytokinin inactivation, genes coding for the N-glucosyltransferase *UGT76C2* (Wang *et al*., 2011), and the genes that encode for enzymes responsible for cytokinin degradation *CKX4* and *7* (Werner *et al*., 2001, 2003; Köllmer *et al*., 2014) presented expression changes upon NTT induction. In general, cytokinin-related genes presented greater changes in expression than auxin-related genes. We decided to analyze *IPT5*, *AHP6* and CKX7 as genes that act at different steps of the cytokinin pathway and could act downstream NTT, because of their reported expression patterns (Miyawaki *et al*., 2004; Atta *et al*., 2009; Köllmer *et al*., 2014) already suggested some coincidences with *NTT* expression. *IPT5* has been reported to be expressed in incipient lateral roots (Miyawaki *et al*., 2004), like *NTT*. Moreover, the transcriptional fusion reporter line *pIPT5::GUS* showed expression in the vasculature of different tissues and the SAM, where *NTT* expression was also detected. The *pAHP6::GFP* reporter line shows expression in the root vasculature (Mähönen *et al.,* 2006; Moreira *et al.,* 2013). Finally, *CKX7* is expressed in the transmitting tract and the vasculature of different organs (Köllmer *et al*., 2014), including the replum (Di Marzo et al., 2020), where *NTT* is also expressed (Chung *et al*., 2013; Marsch-Martínez *et al*., 2014, and Fig. 1 M, O).

### NTT induction causes an increase in *IPT5* expression

Given the increase in cytokinin content detected upon NTT induction, we first focused on studying the cytokinin biosynthesis gene *IPT5*. We analyzed the changes in its expression after NTT induction by RT-qPCR and found an increase in *IPT5* expression as occurred in the transcriptome analysis. The ratio of the increase was also comparable to the ratio found in the transcriptome analyses (Fig. 3A).

**Figure 3.**
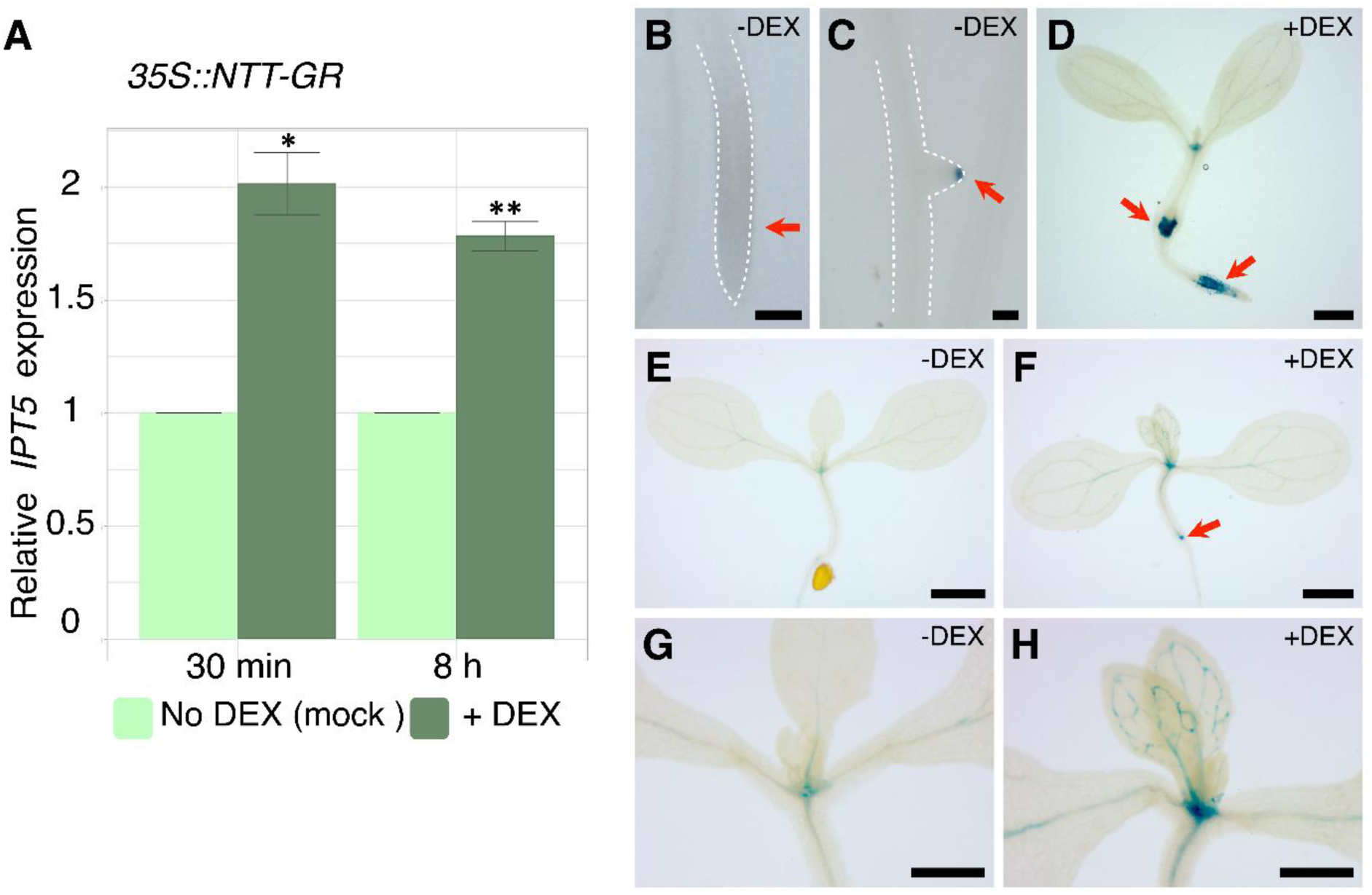
Increase of *IPT5* expression after NTT induction in RT-qPCR and genetic assays. (A) *IPT5* relative expression at 30 min and 8 h after NTT induction in DEX treated (+DEX, dark green) and mock treated (-DEX, light green) root tissue. The y-axis indicates the fold change. Three biological replicates were used for each measurement. Values are given as mean ± SE. Asterisks above the bars indicate significant differences between the respective values (* p<0.05, ** p < 0.01, Student’s t-test). (B-H) *35S::NTT-GR x pIPT5::GUS* seedlings mock treated or induced with DEX and stained 48 h after induction. (B-D) seedlings that were geminated directly on either mock (B, C) or DEX (D), supplemented medium and stained 7 dag. Red arrow indicating absence of staining in main root (B) and staining in the tip of a lateral root (C); (D) seedling germinated directly in media with DEX, red arrows indicate staining in elongation zone and at the hypocotyl-root junction, barely detected in non-induced plants. (E, G) mock treated seedlings, showing staining in SAM and slightly in cotyledon vasculature; (F, H) seedlings induced with DEX. The red arrow indicates GUS staining at the hypocotyl-root junction, similar to D. Also, there is an increase of staining in vasculature. G and H are a close ups from E, F respectively. Scale bars 0.1 mm in (B, C), 1 mm in (E, F), and 0.5 mm in (D, G, H).

We crossed an *IPT5* reporter line (*pIPT*5::*GUS,* Dello Ioio *et al.,* 2007) with the NTT inducible line to evaluate its spatial expression upon NTT induction. In *35S::NTT-GR x pIPT*5::*GUS* seedlings, we observed an increase in GUS staining 48 hours after NTT induction (Fig. 3B-H). In the shoot, the increase in *pIPT*5::*GUS* expression was detected particularly in the SAM and in the vasculature of developing leaves (Fig. 3E-H). Also, we observed GUS staining at the base of the hypocotyl, which was barely detected in non-induced plants (Fig. 3D, F). Moreover, strong GUS staining was detected in the root elongation zone in seedlings that were directly germinated in DEX-supplemented medium, contrasting with no staining in this zone in non-induced seedlings (Fig. 3B-D). Therefore, *IPT5* expression showed a clear increase after NTT induction. This increase appeared to be specific to certain tissues, and more intense in roots.

### NTT affects cytokinin response

After finding that NTT could alter cytokinin content and *IPT5* expression, we then evaluated the effect of NTT on cytokinin signaling. Since *AHP6* was one of the signaling genes with changed expression, we explored its expression after NTT induction both by RT-qPCR and using the *pAHP6::GFP* reporter line (Fig. S4). When root tips of non-induced and induced seedlings were visualized, we observed a decrease in the GFP signal in the induced seedlings (Fig. S4A-D), the same tendency as the transcriptome and RT-qPCR analyses (Fig. S4E). Because the transcriptome analyses revealed that more signaling genes were also affected by NTT induction, we explored the overall effect of NTT induction in the cytokinin pathway in roots, using a cross of the NTT inducible line with the *TCSn::GFP* cytokinin reporter line (Zurcher *et al*., 2013).

We analyzed cytokinin transcriptional response 48 h after NTT induction and observed that in the meristem cell elongation transition zone, *TCSn::GFP* signal was visible in the vascular bundle and epidermis of induced roots but absent in mock-treated roots (Fig. 4A, B). The same occurred in the mature region of the root, where the GFP signal was observed in the vascular bundle and epidermis of induced plants, but barely detected in non-induced plants (Fig. 4C, D). Conversely, at the root apical meristem, though slight differences were observed, the *TCSn::GFP* pattern was similar in mock and DEX treated plants, with a clear signal in the lateral root cap, columella, stele initials, and the cells above them (Fig. 4E, F, Zürcher *et al.,* 2013). However, 7 days after induction, no *TCSn::GFP* signal was visible in the root apical meristem anymore, and meristem organization was altered (Fig. 4G). Nevertheless, in these same roots, but at upper regions, the *TCSn::GFP* signal was still visible in newly formed lateral root primordia (Fig. 4H).

**Figure 4.**
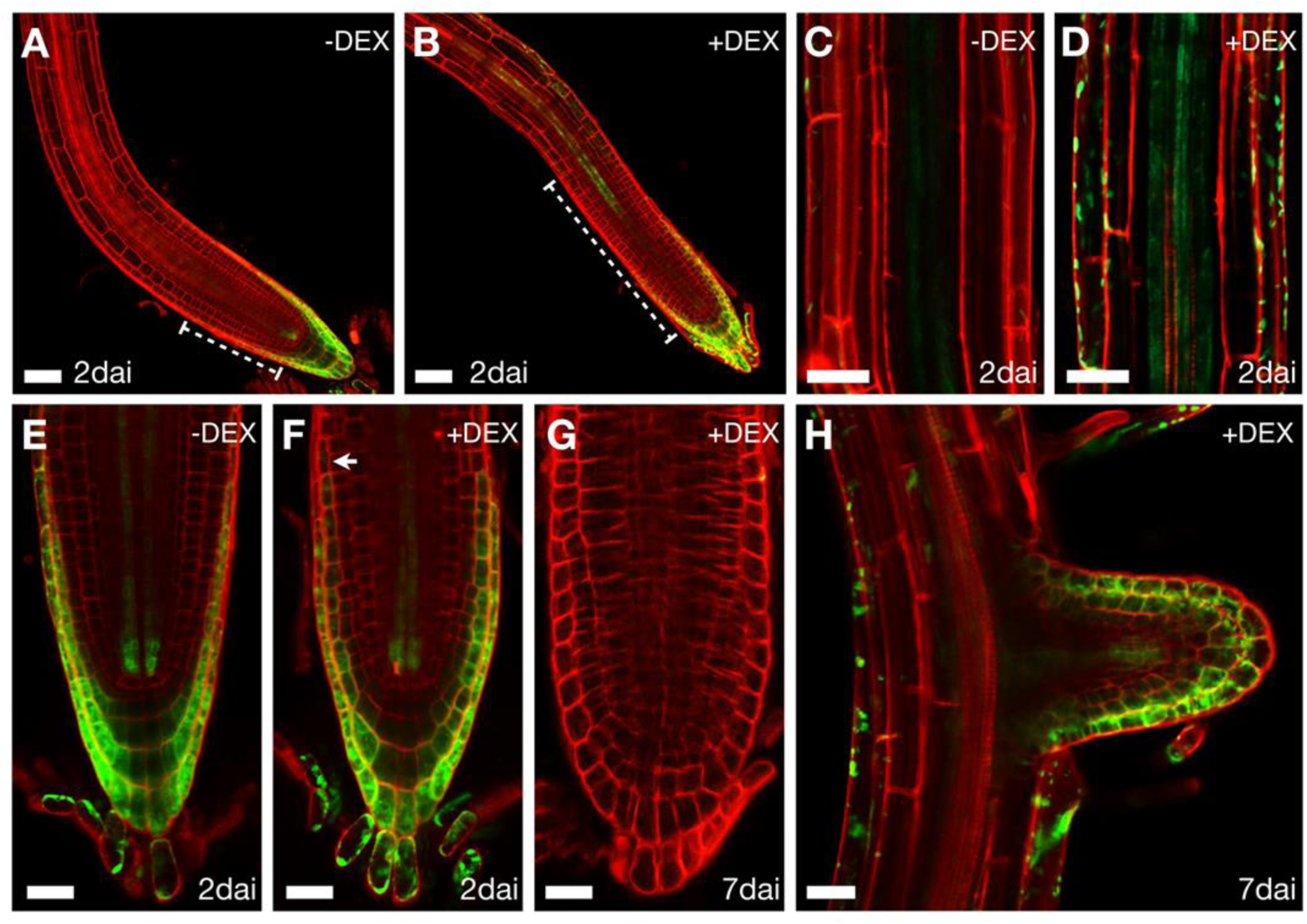
Cytokinin signaling (*TCSn::GFP*) response to NTT induction. (A-H) 4 dag *35S::NTT-GR x TCSn::GF* seedlings either mock or DEX treated, observed 2 dai (A-F) or 7 dai (G-H). (A) In mock treated roots, no signal is observed beyond the meristematic zone; (B) In contrast, expression is detected in the vasculature between the proliferation and transition domains in induced plants 48 h after induction (2 dai). (C, D) In differentiated cells in mature roots, no expression can be seen in mock treated plants (C), while expression is detected in DEX treated seedlings 2 dai (D). (E) Mock treated RAM observed two days after treatment. (F) Changes in *TCSn* expression in DEX treated roots are almost imperceptible in the root tip 2 dai, but an expansion of the signal upwards in the second external cell layer was detected when compared to mock treated roots (white arrow). (G) More evident changes can be noticed 1 week after induction, when the morphology of the RAM seems clearly different, and the signal is lost in DEX treated plants. (H) This same plant still shows expression in young lateral roots. Scale bars in (A, B) represent 50 µm, and in the rest of the panels 25 µm.

Therefore, cytokinin signaling is affected after NTT induction in diverse ways depending on the tissue and time after induction, with some regions increasing and others decreasing CK response.

### *CKX7* mediates part of the NTT induction phenotype

Intriguingly, even though different cytokinin biosynthesis genes were upregulated and CK content was increased, also the expression of cytokinin degradation or inactivation genes increased after NTT induction. Moreover, CK signaling increased or decreased depending on the tissue, suggesting a complex effect of NTT on the CK pathway. *CKX7,* that encodes a cytokinin degradation enzyme, was found among the genes that were differentially expressed. It was upregulated in three of the four conditions and showed the greatest upregulation compared to other auxin- or cytokinin-related genes. When its expression was evaluated by RT-qPCR, a clear increase was again detected (Fig. 5A). We found puzzling that a cytokinin degradation gene was upregulated by NTT along with *IPT5*, and we investigated whether *CKX7* could also be involved in the NTT induction phenotype. For this, we analyzed the effect of the loss of *CKX7* function in the *35S::NTT-GR* phenotype.

**Figure 5.**
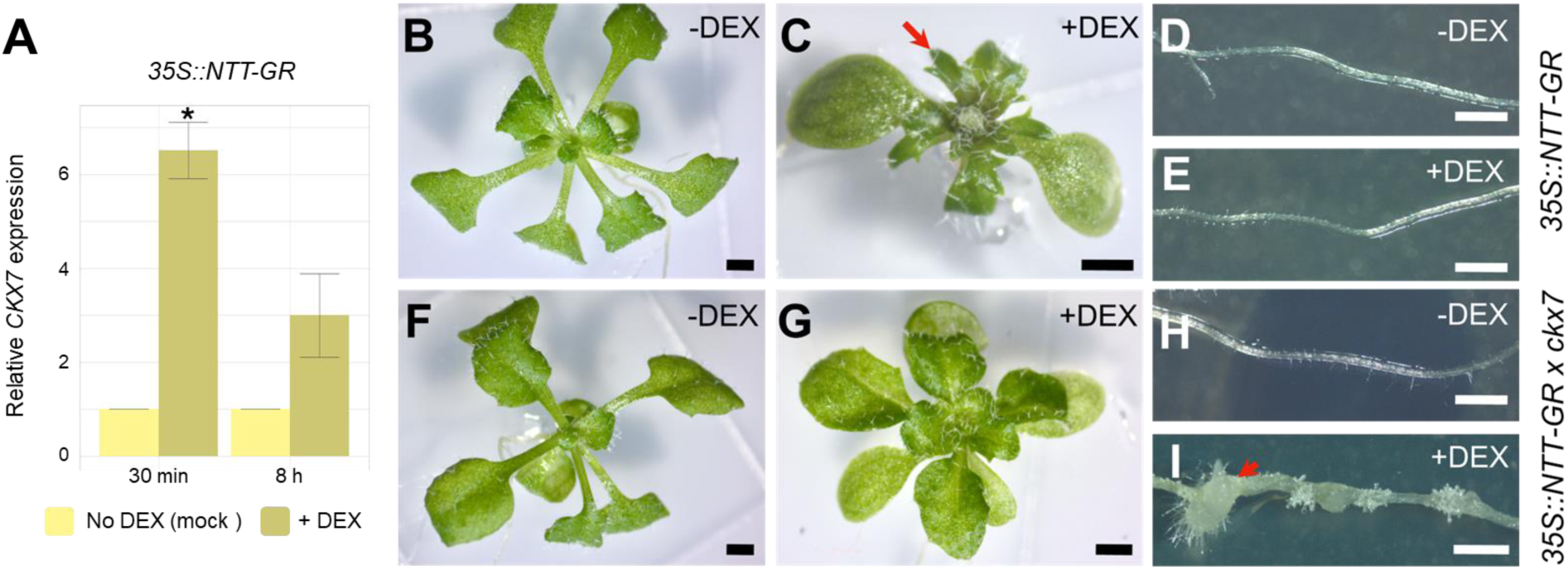
NTT induction phenotype in *ckx7* seedlings. (A) *CKX7* relative expression at 30 min and 8 h after treatment in root tissue treated with DEX (+DEX, dark yellow) or mock-treated (-DEX, light yellow). The y-axis indicates the fold change. Three biological replicates were used for each measurement. Values are shown as mean ± SE. Asterisks above the bars indicate significant differences (*p<0.05, Student’s t - test). (B-I) Mock of DEX treated *35S::NTT-GR* compared to *35S::NTT-GR* x *ckx7* plants. Shoots of *35S::NTT-GR* (B, C) and *35S::NTT-GR* x *ckx7* (F, G) seedlings, were either mock treated (B, F), or induced with DEX (C, G). Roots of *35S::NTT-GR* (D, E) and *35S::NTT-GR* x *ckx7* (H, I) seedlings, either mock treated (D, H), or induced with DEX (E, I). Red arrows highlight irregularities in leaf margins or callus formation in roots. Scale bars represent 1 mm in all images.

Uninduced *35S::NTT-GR* and *35S::NTT-GR ckx7* plants were very similar and resembled wild-type plants (Fig. 5B, F). As expected, DEX-induced *35S::NTT-GR* plants developed leaf abnormalities as shown before (Fig. 1B, 5C). Strikingly, the *ckx7* mutation largely suppressed this effect. The leaves of the plants did not form pronounced leaf projections or serrations when induced, although they still presented shorter petioles (Fig. 5G). Most leaves in the *35S::NTT-GR ckx7* individuals examined after induction did not present serrations at all or only very slight serrations. Therefore, CKX7 activity appears to be required for the irregular leaf margin phenotype caused by NTT induction. Moreover, the leaves of induced *35S::NTT-GR ckx7* were wider and larger than those of induced *35S::NTT-GR* (Fig. 5C, G). Interestingly, induced *35S::NTT-GR ckx7* plants presented an unexpected root phenotype, not observed in induced *35S::NTT-GR* plants (Fig. 5D, E, H, I). DEX-treated *35S::NTT-GR ckx7* roots developed white calli, clearly visible 16 days after induction, at the positions where secondary roots would usually emerge (Fig. 5 I). Therefore, *CKX7* prevents the formation of calli in the roots of induced plants, probably by locally counteracting the increased CK content.

We also tested the effect of adding cytokinins (BAP) to both *35S::NTT-GR* and *35S::NTT-GR ckx7* plants. Uninduced *35S::NTT-GR* that were treated with BAP produced serrated leaves with short petioles and a profusion of trichomes (Fig. S6A). However, uninduced *35S::NTT-GR ckx7* were differently affected by the cytokinin treatment. Most of their leaves also presented serrations and shorter petioles, but they were evidently smaller, and some seedlings presented one larger leaf (Fig. S6E), suggesting that *35S::NTT-GR ckx7* seedlings display an increased response to the supplemented cytokinin, as would be expected. When *35S::NTT-GR* seedlings grown on cytokinin-supplemented medium were induced by DEX, they developed much smaller leaves, shorter petioles, and more severe leaf serrations or projections at the leaf margin, than those grown on normal medium (Fig. S6A, B). *35S::NTT-GR ckx7* plants under the same conditions also developed leaves with very short petioles (Fig. S6F), but these leaves were evidently wider and larger than those of cytokinin treated *35S::NTT-GR* plants. Instead of serrations, their edges presented undulations, leading to an atypical shape of lettuce-like appearance (Fig. S6F). The undulations and trichomes were more evident (Fig. S6F) than those of induced *35S::NTT-GR ckx7* plants without cytokinins (Fig. 6G).

**Figure 6.**
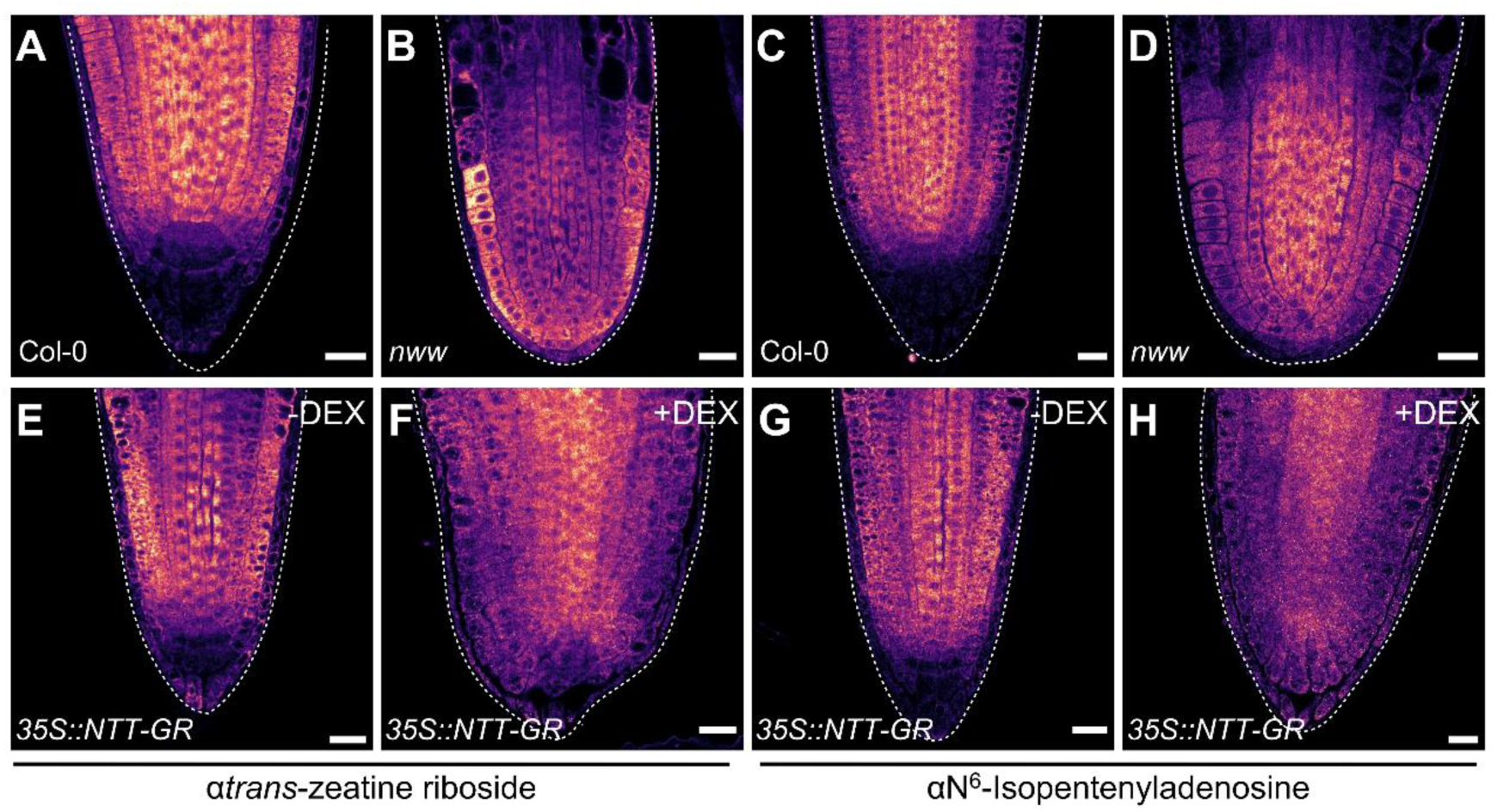
Cytokinin immunolocalization in root tips. Whole-mount root tips of 1 dag Col-0 (A, C), *nww* (B, D) and *35S::NTT-GR* (E-H) seedlings, either mock treated (E, G), or induced with DEX (F, H). LUT images of the localization of trans-Zeatin-riboside (A, B, E, F), or Isopentenyl adenosine (C, D, G and H). Scale bars represent 20 µm.

None of the uninduced *35S::NTT-GR ckx7, 35S::NTT-GR,* or wild-type plants produced calli when treated with cytokinins (Fig. S6C, G). When induced *35S::NTT-*GR seedlings were treated with cytokinin, only one in 40 plants developed calli (Fig. S6D), in contrast to 40 out of 40 induced, cytokinin-treated, *35S::NTT-GR ckx7* plants (Fig. S6H).

Taken together, the results indicate that *CKX7* acts downstream of NTT and plays a significant role in the phenotypes that NTT induction causes in the plants, possibly by modulating the cytokinin content locally.

### The CK localization pattern is altered in *nww* mutants and after NTT induction

After observing the changes in overall cytokinin content and the relevance of *CKX7* in the NTT induction phenotype, which would affect local cytokinin content, we sought to explore cytokinin localization in the mutant and the inducible overexpression backgrounds. *NTT* expression is localized to a few cells in the meristem of the young developing root (Fig 1L), where it plays an essential but redundant function with *WIP4* and *WIP5* (Crawford *et al*., 2015). Therefore, the triple mutant lacks roots (Crawford *et al*., 2015), which can be distinguished in very young seedlings. Therefore, we analyzed the cytokinin localization pattern by immunolocalization at the root tip in young wild-type and inducible overexpressors, or the equivalent region in the *nww* mutant (Fig. 6).

Trans-zeatin and isopentenyl adenine content was changed in roots after NTT induction. Moreover, trans-zeatin content is reduced to only 9% of the normal wild-type content in plants that overexpress *CKX7* (Kollmer *et al*., 2014). Therefore, we used anti-trans-zeatin riboside and anti-N6-isopentenyladenosine antibodies to assess their localization. Interestingly, the pattern of cytokinin distribution in the root tip was clearly affected both in the mutants and the induced lines. Wild-type (Fig. 6A, C) and *35S::NTT-GR* non-induced (Fig. 6E, G) roots had a similar pattern of cytokinin distribution. In both, trans-zeatin riboside and isopentenyl adenosine were abundant above the quiescent center of the meristem and relatively low below it and in the columella. In the *nww* mutant, however, they were more homogeneously distributed towards the apex, and reduced towards the top (Fig. 6B, D), compared to wild-type roots (Fig. 6A, C). In the case of the induced *35S::NTT-GR* plants, they presented a homogeneous distribution in the whole root tip, which already displayed an altered size and morphology (Fig. 6 F, H), compared to non-induced plants (Fig. 6 E, G).

In conclusion, both the mutant and the inducible overexpressor line showed clear alterations in the distribution of the immunodetected cytokinin types.

### NTT binds to the promoter of cytokinin-related genes

The effects of NTT in cytokinin homeostasis and the expression of cytokinin-related genes were very clear. Because the translation inhibitor cycloheximide was used in the transcriptome and RT-qPCR analyses at 30 min, some differentially expressed genes could be direct NTT targets. We tested whether the regulatory regions of the three selected genes (*IPT5*, *AHP6* and *CKX7*) were bound by NTT, first using Y1H assays.

We sought predicted NTT binding sites in the promoters and coding regions (Persikov and Singh, 2014; Herrera-Ubaldo *et al.,* 2019) of these genes and cloned genomic fragments containing them (Fig. 7A). We also tested other promoter fragments. For *IPT5*, we tested a 1945 bp promoter, a 70 bp promoter fragment (*IPT*5-A), and a 435 bp fragment inside the coding region that contains two predicted binding sites (*IPT*5-B). The *AHP6* promoter presented a single site, 720 bp upstream of the ATG. We tested a 1900 bp promoter fragment, and a 450 fragment around the predicted site (*AHP6*-A). For the *CKX7* regulatory region, we used a fragment comprising 969 bp upstream the ATG, containing two predicted sites (Fig. 7A).

**Figure 7.**
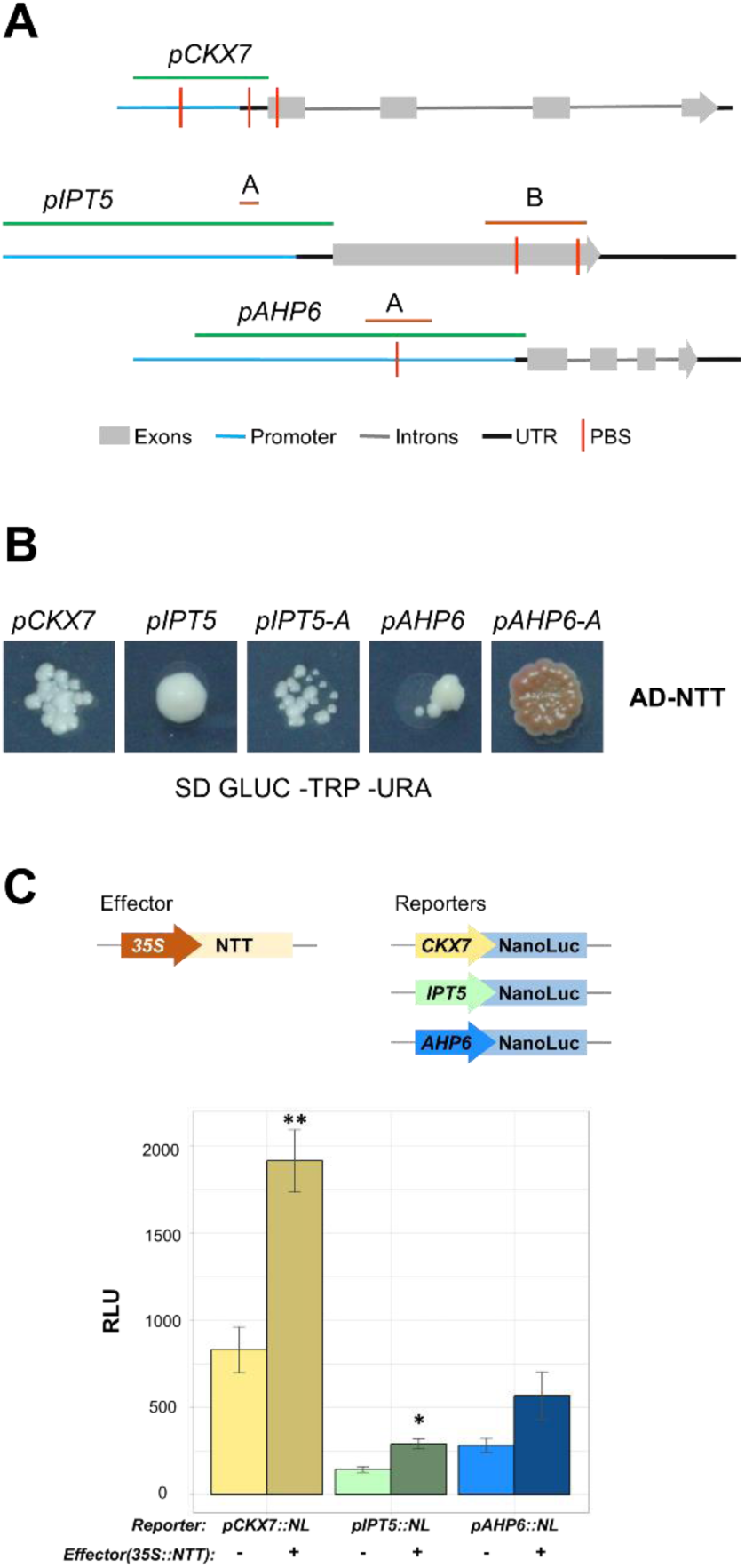
Y1H and NanoLuc assays to test NTT binding and regulation of candidate target genes. (A) Schematic representation of the tested genomic regions, with large promoter fragments shown in green and short or internal fragments in orange. PBS: Putative Binding Site. (B) Y1H analyses of NTT protein interaction with the target promoters and fragments depicted in (A). (C) NanoLuc assays. The constructs used as effector and reporters are indicated in the diagrams. The graph presents relative NanoLuc activity, detected in each combination. Values are given as mean ± SE and asterisks indicate a significant difference (* p ≤ 0.01, ** p ≤ 0.001, Student’s t-test, RLU: Relative Luminescence Units).

We found that NTT binds the three complete promoters of *IPT5, CKX7* and *AHP6* (Fig. 7B). NTT also bound to the short *IPT*5-A fragment (Fig. S5). The other fragment, *IPT5-B*, could not be tested due to autoactivation (Fig. S5). Moreover, NTT bound to the *AHP6-A* fragment (*AHP6*-A, Fig. 7B). Therefore, it appears that NTT could be directly regulating the expression of *CKX7, IPT5* and *AHP6.* To further confirm other differentially expressed genes, we also evaluated NTT binding to the promoters of *KAN*, which was downregulated in all samples, and *PUP10*, a cytokinin transporter downregulated in roots at both early and late times. NTT also bound to these promoters in the Y1H test (Fig. S5).

Next, we validated and explored the regulation of NTT on *IPT5, CKX7* and *AHP6* promoters *in planta* using a transient luciferase assay. The same complete *IPT5, CKX7* and *AHP6* promoters used for the Y1H assay were fused to NanoLuc (Urquiza-García and Millar, 2019; Becerra-García *et al*., 2023). These promoters were transiently expressed in tobacco leaves, in combination with a *35S::NTT* construct (Fig. 7C). For *pIPT5* and *pCKX7*, a clear activation of expression was revealed by a marked increase in bioluminescence when *35S::NTT* was co-expressed with the promoter constructs (Fig. 7C). This indicates that NTT positively regulates these promoters in this system. These genes were found to increase their expression in the transcriptome data and by RT-qPCR 30 min after NTT induction, suggesting that they may be target genes positively regulated by NTT. In the case of the *AHP6* promoter, although there was an increase in NanoLuc activity when coexpressed with NTT, it was not statistically significant, This gene was downregulated in the root, in the transcriptome analyses and RT-qPCRs performed in induced seedlings and may be downregulated or upregulated in other tissues and stages.

Together, the results of this work indicate that NTT can modulate the cytokinin pathway at different steps in a complex fashion.

## Discussion

In this work, we investigated whether the NTT/WIP2 transcription factor regulates genes involved in the cytokinin pathway. Some phenotypes of NTT overexpression lines are similar to those of plants that have been grown in cytokinin-supplemented medium (Chung *et al*., 2013; Marsch-Martínez *et al*., 2014; Skalák *et al*., 2019; Dello Ioio *et al*., 2007; Liu *et al*., 2022; Werner *et al.,* 2003). Also, cytokinin treatments can recover replum development in the Arabidopsis *rpl ntt* double mutant, which lacks this tissue (Zuñiga-Mayo *et al*., 2018). Moreover, STK, an interacting partner of NTT (Herrera-Ubaldo *et al*., 2019) regulates cytokinin levels during fruit development (Di Marzo *et al.,* 2020), and NTT/WIP2 has been found to interact with cytokinin signaling system components (Dortay *et al*., 2008). In addition, NTT/WIP2 is a target of ARF5/MP, thought to act downstream auxin signaling (Crawford *et al*., 2015), and the auxin and cytokinin pathways are frequently closely related (Schaller *et al*., 2015). When revising published expression patters of the cytokinin response TCS marker (Muller and Sheen, 2008), there seemed to be coincidences with reported NTT expression during very early stages of root development (Crawford *et al*., 2015), the development of the gynoecium, particularly in the inner tissues (Crawford *et al*., 2007; Marsch-Martinez *et al*., 2012), and the vasculature (as revealed by the *TCSn::GFP* line; Zürcher *et al*., 2013; Skalák *et al*., 2019).

Accordingly, changes in cytokinin content were found in seedlings where the activity of NTT was induced. Curiously, the changes in content were different depending on the tissue (roots or shoots), suggesting that NTT could have a distinct, tissue-specific effect on cytokinin content.

### NTT affects cytokinin genes and has a tissue-specific effect on gene expression

Global expression analyses provided information about candidate cytokinin genes that could be modulated by NTT. An inducible NTT line was used because of the high degree of redundancy of NTT with other WIP family members (Du *et al*., 2022) and the severe phenotype of combined mutations (Crawford *et al*., 2015). Interestingly, the differentially expressed cytokinin genes participate in a positive (such as the biosynthetic enzymes IPT5 or LOG4 or the type-B ARRs) or a negative way (such as cytokinin degradation enzymes CKX7, or the negative signaling components AHP6, or type-A ARRs) in cytokinin metabolism, signaling or response. Moreover, the changes in expression of cytokinin-related genes were higher than for genes belonging to the auxin pathway. This suggests that NTT could modulate the cytokinin pathway at diverse steps and possibly have a stronger effect in it than in the auxin pathway at a global level.

Regarding the overall data obtained from the RNA-seq experiments, many interesting DEGs were found and worth of further analyses in future studies, like *CUC1* and 2, *KNAT1, 4, 6* and *7, AS2, KAN, WOX4, SHR* and *NGA3*.

The data obtained from roots and shoots was analyzed separately, and some DEGs were found in both tissues at both times, suggesting that they may be directly or indirectly regulated by NTT independently of the tissue. One example of these common DEGs was *KAN* (Fig. S3B, Bowman *et al.,* 2001). Its promoter is bound by NTT in Y1H experiments (Fig. S5) and was also found to be downregulated and a direct target of NTT (Gomez-Roldán *et al*., 2020) and the closely related WIP5 (O’Malley *et al.,* 2016). However, we also found DEGs, the majority, for which the change in expression was dependent on the tissue and time after induction. Therefore, NTT might have tissue-specific activity and affect genes differently depending on the context. In the GO category enrichment analyses, again, both common and tissue specific categories were present, and some were similar to those found in other reports. For example, in *Gerbera hybrida* 45-day-old flower bud clusters that overexpress *GhWIP2,* an *NTT* homologue, *GhWIP2* was reported to act as a repressor, suppressing cell expansion during organ growth by modulating crosstalk between gibberellins, abscisic acid, and auxin (Ren *et al.,* 2018). Categories related to these hormones were also enriched in our transcriptome comparisons.

WIP1 and 2 proteins have been reported to integrate hormonal signals to regulate organ growth and adaptation to stress (Gomez-Roldan *et al*., 2020). Most cell wall, development, growth, and morphogenesis categories were enriched among downregulated genes in whole seedling samples of inducible WIP1 and WIP2/NTT lines analyzed by Gomez-Roldán *et al*., 2020. In contrast, when the tissues were analyzed separately (this work), cell wall, development and morphogenesis categories were prominently present in the root 8 h downregulated genes but were less prominent in aerial tissue DEGs. Stress categories, enriched among the upregulated genes in whole seedlings (Gomez-Roldán *et al*., 2020), were also frequent in both tissues (this work). However, while genes belonging to general hormone categories were present in all conditions, those belonging to specific hormonal categories, such as jasmonic acid, ethylene, gibberellins, cytokinins and abscisic acid, are enriched differently in the distinct tissues and times (Dataset S2).

Therefore, NTT may regulate genes and processes both in a general and a tissue-specific (young shoot or root, as observed for changes in cytokinin content), or maybe even stage-specific manner. This may also occur in other organs and tissues, such as the gynoecium, the vasculature, or specific cells at certain stages, like during early embryo development.

### NTT modulates cytokinin homeostasis

Since our aim was to study the effect of NTT on the cytokinin pathway, we focused on cytokinin genes, particularly those with reported expression in tissues where NTT is also expressed, such as transmitting tract, vasculature and young roots, and where cytokinin homeostasis plays an important role (Miyawaki *et al*., 2004; Atta *et al*., 2009; Köllmer *et al*., 2014; Miyawaki *et al*., 2004; Mähönen *et al.,* 2006; Moreira *et al.,* 2013; Köllmer *et al*., 2014; Di-Marzo *et al*., 2020; Muller and Sheen, 2008; Chung *et al*., 2013; Marsch-Martínez *et al*., 2012 and 2014; Zurcher *et al.,* 2013).

Cytokinin levels are the result of the balance between biosynthesis and degradation or inactivation pathways (Kieber and Schaller, 2018), and transport impacting their distribution among different tissues (Hirose *et al*., 2008). For biosynthesis, the rate-limiting step is catalyzed by ADP/ATP isopentenyl transferases (IPTs, Miyawaki *et al*., 2004). *IPT* increased expression results in higher levels of iP (Kakimoto, 2001; Sun *et al*., 2003; Zubko *et al*., 2002), while in *ipt* mutants, iP and tZ-type cytokinin species are reduced (Miyawaki *et al*., 2006). *IPT5* was increased upon NTT induction, and accordingly, a higher content of total, and specially iP, tZ and DHZ (a reduced form of tZ) types, was detected in the roots of induced plants. Interestingly, even though the inducible construct contains a 35S promoter, *pIPT5::GUS* expression increased only in specific regions of the plant, suggesting that *IPT5*, and possibly many other target genes, are subject to tissue specific regulation.

The expression pattern of the cytokinin response marker TCS was also affected by NTT induction. Cytokinins are perceived by membrane receptors which start a signaling phosphorelay cascade that activates type-B ARR transcription factors (Kieber and Schaller, 2018). The TCS signal both increased and appeared in ectopic regions or decreased and disappeared from its normal regions of expression, depending on the region of the tissue, root in our case, and time after induction, again suggesting cytokinin homeostasis is affected after NTT induction in different ways depending on the tissue and time after induction. The TCS marker reports the transcriptional response to cytokinin, reflecting both content and signaling activity (Muller and Sheen, 2008; Zürcher *et al*., 2013). Therefore, the observed changes could be due to expression differences in genes acting at different steps of the pathway, as found in the transcriptome comparisons. One of these genes is *AHP6*, which plays an important inhibitory role in the signaling cascade (Mähönen *et al.,* 2006), and its expression was confirmed to be altered upon NTT induction.

Cytokinins could be synthesized locally, transported, act as an autocrine or paracrine signals, and be catabolized at nearby or distant sites (Hirose *et al*., 2008). Moreover, cytokinins can have opposite roles depending on the developmental stage of an organ or tissue (Skalák *et al*., 2019; Di Marzo *et al*., 2020). Therefore, regions where cytokinin was crucial, may require cytokinin degradation to further proceed in their development. For degradation, the cytokinin catabolic enzymes CKX are key in controlling cytokinin levels in plant tissues (Werner *et al*., 2001). *CKX7* presented a high change in expression in the global expression analyses, RT-qPCRs and transactivation assays. From the members of the Arabidopsis CKX family, it is the only localized at the cytosol (Werner *et al*., 2001, 2003; Köllmer *et al.,*2014) and has been proposed to have unique access to a specific subcellular pool of cytokinin, relevant in tissues such as the root procambium (Köllmer *et al.,* 2014). Its overexpression causes root inhibition and a higher reduction on *c*Z-type cytokinins (*c*Z and *c*Z9G) and iP9G, in comparison to CKX1 and CKX2 (Köllmer *et al.,* 2014). Interestingly, after NTT induction, while the content of most cytokinin types increased in induced roots, cZ species were not, and iP, iP7G and iP9G content was even reduced in shoots. In Arabidopsis, *CKX7* is expressed in the transmitting tract and root and shoot vasculature (Köllmer *et al*., 2014), two regions where *NTT* is also expressed. Moreover, *CKX7* has been recently reported to be a target of STK, an NTT protein interactor (Di Marzo *et al*., 2020; Herrera-Ubaldo *et al*., 2019), suggesting that *CKX7* could be a common downstream target gene of both transcription factors. The expression found in the cycloheximide-treated samples of the transcriptome, RT-qPCR, transactivation, and Y1H assays also support this.

Interestingly, the lack of *CKX7* function in the NTT inducible line led to a clear reduction in the induction phenotype, indicating that at least partially, NTT acts through *CKX7*. The reduction of serrations was unexpected, and we speculate that the local degradation of cytokinins by CKX7 contributes to patterns of cytokinin accumulation that are involved in the formation of serrations or morphological changes in leaf edges. Moreover, the appearance of calli in NTT induced *ckx7* roots could be related to the reduced capacity of the plants to counteract the cytokinin excess promoted by NTT, in the absence of *CKX7* function.

### NTT modulates the localization pattern of cytokinins

The auxin-cytokinin interplay has been compared to the yin-yang concept of complementarity (Schaller *et al*., 2015). They are connected both positively and negatively and together guide the development of the early embryo (Muller and Sheen, 2008), the vasculature in roots (Mähönen *et al.,* 2006; De Rybel *et al.,* 2014; Ohashi-Ito *et al.,* 2014), or the gynoecium (Marsch-Martinez *et al.,* 2014; Zuñiga-Mayo *et al.,* 2014), among others. We found that NTT, an ARF5/MP target (Crawford *et al*., 2015), regulates cytokinin related genes, suggesting that it can link these two pathways (Fig. 8). Callus development is another process in which both auxin and cytokinin play a crucial role (Skoog and Miller, 1957). The formation of calli in the induced *35S::NTT-GR ckx7* line could point to the connection between both pathways. Moreover, interestingly, in ovules, a different ARF5/MP isoform is expressed in cells with low auxin and high cytokinin concentration (Cucinotta *et al*., 2021), suggesting that MP can be related to both.

**Figure 8.**
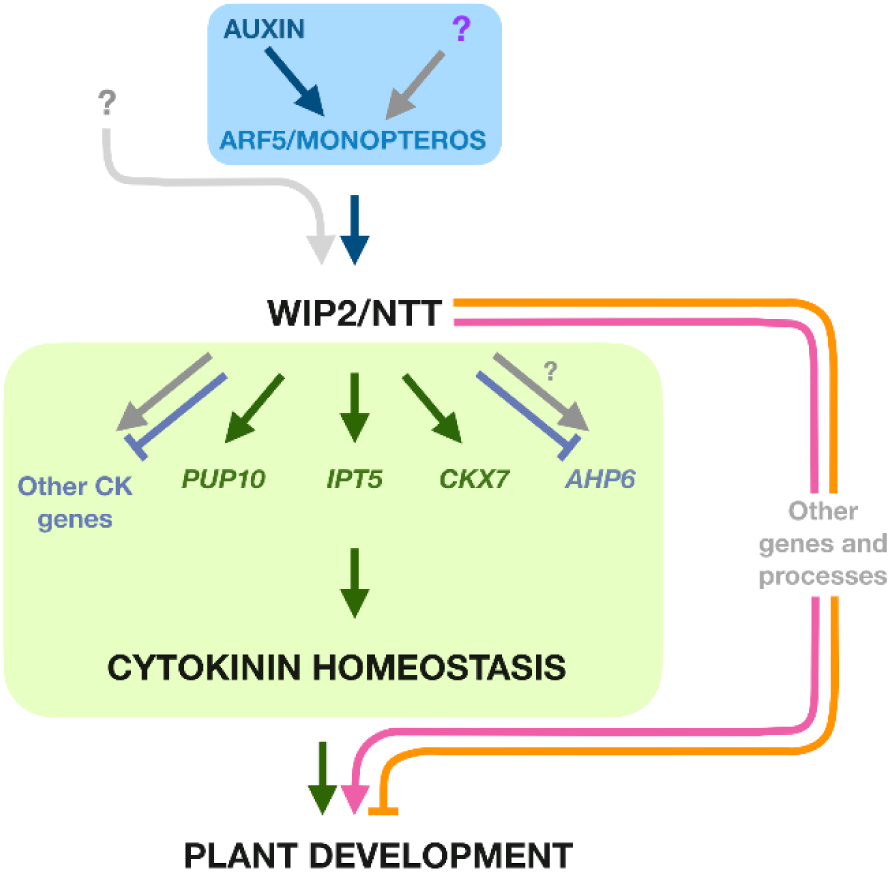
A model of NTT regulation. Proposed model of NTT acting as a link between the auxin and the cytokinin pathways. Additional stimuli beyond auxin and ARF5/MP may regulate *NTT* expression. NTT modulates the expression of genes involved in cytokinin biosynthesis, signaling, and degradation, among others.

Another MP target, *TARGET OF MONOPTEROS 5* (*TMO5*), with *TMO5-LIKE1* (*T5L*) and their interacting partner LONESOME HIGHWAY (LHW), activate the expression of the cytokinin genes *LOG4* and *AHP6* (Ohashi-Ito *et al.,* 2007, 2014; De Rybel *et al*., 2014). These genes, together with *CKX3*, lead to the creation of a local, nonresponding cytokinin source within the root vascular tissue that is essential for its proper development (Ohashi-Ito *et al*., 2014; De Rybel *et al.,* 2015; Yang *et al*., 2021).

We speculate that a similar situation, albeit through the regulation of different genes, could occur for NTT. *CKX7* upregulation, together with *IPT5* and *AHP6* modulation, if simultaneous, may generate cytokinin differential regions, either in content or in response within the NTT-responsive regions. CKXs have been proposed to participate in the maintenance of cytokinin local developmental fields (Werner *et al*., 2001). Moreover, Köllmer *et al.,* 2014 noticed that both *pCKX7* and the cytokinin reporter *TCSn::GFP* (Zürcher *et al.,* 2013) are expressed in an unequal distribution in the embryo sac and already proposed that CKX7 could establish a cytokinin gradient in the cytoplasm (Köllmer *et al.,* 2014). It would be interesting to explore these possibilities.

Another hypothetical scenario to explain the activation of both *IPT5* and *CKX7* by NTT could be a short temporal delay between their activation. Interestingly, some of the tissues where *TCSn::GFP* and *NTT* coincide present early cytokinin signaling that is lost at later stages, like at the medial domain of the gynoecium (Marsch-Martinez *et al*., 2014; Reyes-Olalde *et al.,* 2017), or the first division of the hypophysis in the embryo, after which one cell retains, and one loses TCS signal (Muller and Sheen, 2008). *NTT* is expressed in these two tissues (Crawford *et al*., 2007, 2015) and could first activate *IPT5* to increase cytokinin content, and then *CKX7* to decrease it when required.

Nevertheless, even if the mechanism by which the regulation of the cytokinin pathway by NTT works is not yet clear, it is evident that NTT impacts cytokinin distribution. The immunolocalization assays performed for two cytokinin species in the *nww* mutant and the inducible line, showed clear changes in their distribution pattern.

The distribution of specific CK species is the result of production in certain tissues, transport, and modification in the same or in other tissues, as CK species are interconverted into one another (Mok and Mok, 2001; Hirose *et al*., 2008), and the distribution of active forms will impact the cell transcriptional response (Kieber and Schaller, 2018). Though *IPT5* and *CKX7* may play a prominent role, it is possible that *PUP10* (whose promoter is bound by NTT), *LOG4* and *7, AHP6, ARRs, KMD1*, and *WOL/AHK4,* genes that were differentially expressed upon NTT induction, among others, could also be involved in the effect of NTT in the cytokinin distribution and transcriptional response. We speculate that the action of NTT in the tissues where it naturally acts, might be also dependent on the tissue and its developmental stage, such as the cells that will give rise to the root meristem, the transmitting tract, and possibly and vasculature, for which a role has not been described, but expression has been observed (Chung *et al*., 2013; Diaz-Ramirez *et al*., 2022). In this regard, *CKX7, AHP6,* or *AHK4/WOL, KAN*, have been reported to also be expressed in vascular tissues or the pericycle (Mähönen *et al.,* 2000, 2006; Emery *et al.,* 2003; Eshed *et al.,* 2004; Hawker and Bowman, 2004; Köllmer *et al.,* 2014). *IPT5* was also expressed in the vasculature. Moreover, the fact that the *nww* mutant is defective in hypocotyl vasculature formation (Crawford *et al.,* 2015) may support this hypothesis.

In conclusion, based on cytokinin quantification, global expression, binding, activation and genetic analyses, together with cytokinin immunolocalization in the mutant and overexpression background, we propose that NTT regulates cytokinin genes and can act as a link between the auxin and cytokinin pathways (Fig. 8). It will be very interesting to further investigate the tissues and developmental stages where this modulation occurs, and whether it results in the generation of cytokinin fields, delayed gene activation or other mechanisms that guide development. Finally, we provide new shoot and root transcriptomics data at different time points, which indicate that NTT modulates a variety of processes, some in a general, and some in a tissue and time-specific fashion.

## Declarations

### Author’s contributions

DDR and NMM conceptualized the research. DDR, NMM, REBG, HHU, YDM, EDA and JECV designed the experiments. ECV, REBG, DDR, HHU, and YDM performed the experiments. RCM performed Bioinformatic analyses. EDA contributed with confocal imaging. ON, MS, MDM, RS, LC and SdF contributed experimental resources and analyzed data. DDR and NMM contributed to manuscript writing and reviewing. NMM obtained financial resources and supervised the project. All authors read and approved the manuscript.

## Acknowledgements

We thank Brian Crawford for plant material and Andrew Millar for NanoLuc plasmids. We thank Andrea Gomez Felipe and Vincent Cerbantez Bueno for sharing some promoter fragments and Juan Carlos Ochoa for his help with RT-qPCR experiments.

## SUPPLEMENTARY DATA

### Supplementary Figures

**Figure S1. Root phenotype of induced *35S::NTT-GR* seedlings.**

**Figure S2. Global expression experiment workflow.**

**Figure S3. Bubble plot of GO enriched categories and heatmap of developmental and auxin-related differentially expressed genes.**

**Figure S4. Change in expression of *AHP6* after NTT induction.**

**Figure S5. Y1H Autoactivation assays, and *pKAN* and *pPUP10* experiments.**

**Figure S6. NTT induction phenotype in *ckx7* seedlings, treated with cytokinin.**

**Figure S7. Root phenotypes and immunolocalization technical control.**

### Supplementary Tables

**Table S1 Primers used in this work.**

**Table S2 Cytokinin profile in shoot and root of Col-0 and *35S::NTT-GR* line, under Mock or DEX treatment (pmol/g FW; Mean ±SE).**

### Supplementary Datasets

**Dataset S1. Differentially Expressed Genes found in the Global Expression Analyses in different tissues.**

**Dataset S2. Gene Ontology enrichment analyses.**

