## Supplemental Table S2 for "The transcription factor NO TRANSMITTING TRACT / WIP2 regulates cytokinin homeostasis in Arabidopsis"

**Table S2 - Cytokinin profile in shoot and root of Col-0 and *35S::NTT-GR* line, under Mock or DEX treatment (pmol/g FW; Mean ±SE).** Cytokinin(CK) types are grouped as follows: Total isopentenyladenine types (Total IP-types), isopentenyladenine (IP), isopentenyladenine riboside (IPR), isopentenyladenine riboside-5′- monophosphate (IPRMP), isopentenyladenine 7-glucoside (IP7G), isopentenyladenine 9-glucoside 50 (IP9G); Total trans-Zeatin (total tZ types): trans-Zeatin (tZ), trans-Zeatin riboside (tZR), trans-Zeatin riboside-5′-monophosphate (tZRMP), trans-Zeatin-O-glucoside (tZOG), trans-Zeatin riboside-O-glucoside (tZROG), trans-Zeatin-7-glucosides (tZ7G), and trans-Zeatin-9-glucoside (tZ9G);Total cis-Zeatin (total cZ types): cis-Zeatin (cZ), cis-Zeatin riboside (cZR), cis-Zeatin riboside-5′-monophosphate (cZRMP), cis-Zeatin-O-glucoside (cZOG), cis-Zeatin riboside-O-glucoside (cZROG), cis-Zeatin-7-glucosides (cZ7G), and cis-Zeatin-9-glucoside (cZ9G); Total dihydrozeatin (DHZ); dihydrozeatin riboside (DHZR), dihydrozeatin riboside-5′-monophosphate (DHZRMP), dihydrozeatin O-glucosides (DHZOG); dihydrozeatin riboside-O-glucoside (DHZROG), dihydrozeatin 7-glucoside (DHZ7G), dihydrozeatin 9-glucoside (DHZ9G). <LOD: not detected. Asterisks indicate statistically significant difference in *35S::NTT-GR* line DEX versus Col-0 DEX (Student's t-test: *p < 0.05, **p < 0.01, *** p < 0.001)

### Cytokinins:

| Tissue | Treatment | Genotype | Total Cytokinins | SE | CK Bases | SE | CK Ribosides | SE | CK Nucleotides | SE | CK O-glucosides | SE | CK N-glucosides | SE |
|---|---|---|---|---|---|---|---|---|---|---|---|---|---|---|
| Shoot | Mock | Col-0 | 96.711 | ±7.55 | 2.288 | ±0.11 | 2.438 | ±0.09 | 32.994 | ±4.78 | 6.860 | ±0.25 | 52.132 | ±2.64 |
| | | *35S::NTT-GR* | 79.618 | ±4.20 | 1.760 | ±0.08 | 1.913 | ±0.26 | 19.809 | ±2.03 | 7.347 | ±0.44 | 48.788 | ±2.67 |
| | DEX | Col-0 | 84.737 | ±3.16 | 4.239 | ±0.59 | 2.109 | ±0.26 | 20.827 | ±2.23 | 7.722 | ±0.39 | 49.840 | ±2.47 |
| | | *35S::NTT-GR* | 87.059 | ±4.80 | 3.929 | ±0.49 | 3.031 | ±0.36 | 20.930 | ±1.59 | 10.235 | ±0.89 | 48.935 | ±3.05 |
| Root | Mock | Col-0 | 337.769 | ±31.78 | 8.802 | ±0.74 | 19.973 | ±2.18 | 37.864 | ±4.97 | 51.310 | ±7.01 | 219.820 | ±18.43 |
| | | *35S::NTT-GR* | 306.268 | ±37.68 | 6.963 | ±0.75 | 16.611 | ±1.92 | 38.327 | ±4.75 | 62.419 | ±8.78 | 181.948 | ±26.02 |
| | DEX | Col-0 | 262.980 | ±23.21 | 6.861 | ±0.34 | 13.661 | ±1.29 | 13.661 | ±2.09 | 43.695 | ±6.17 | 167.471 | ±17.40 |
| | | *35S::NTT-GR* | 516.027 ** | ±50.58 | 8.796 | ±0.81 | 26.543** | ±2.44 | 60.334*** | ±3.24 | 84.057** | ±7.11 | 336.298* | ±39.56 |

### Isopentenyl-types:

| Tissue | Treatment | Genotype | Total iP-types | SE | iP | SE | iPR | SE | iPRMP | SE | iP7G | SE | iP9G | SE |
|---|---|---|---|---|---|---|---|---|---|---|---|---|---|---|
| Shoot | Mock | Col-0 | 50.125 | ±5.71 | 1.014 | ±0.16 | 1.101 | ±0.06 | 23.129 | ±3.36 | 21.579 | ±2.16 | 3.302 | ±0.45 |
| | | *35S::NTT-GR* | 32.956 | ±1.90 | 0.718 | ±0.03 | 0.719 | ±0.09 | 10.232 | ±1.13 | 18.583 | ±0.92 | 2.704 | ±0.16 |
| | DEX | Col-0 | 34.037 | ±1.42 | 0.881 | ±0.10 | 0.879 | ±0.10 | 10.732 | ±1.21 | 18.701 | ±1.00 | 2.845 | ±0.15 |
| | | *35S::NTT-GR* | 24.260* | ±1.87 | 0.766 | ±0.02 | 0.890 | ±0.13 | 7.927 | ±0.40 | 13.001* | ±1.52 | 1.867* | ±0.17 |
| Root | Mock | Col-0 | 44.511 | ±4.14 | 2.491 | ±0.37 | 4.824 | ±0.58 | 8.107 | ±0.75 | 24.717 | ±2.72 | 4.371 | ±0.32 |
| | | *35S::NTT-GR* | 38.146 | ±4.07 | 2.122 | ±0.31 | 2.783 | ±0.36 | 7.116 | ±0.83 | 22.018 | ±2.55 | 4.107 | ±0.62 |
| | DEX | Col-0 | 39.352 | ±3.05 | 2.260 | ±0.28 | 3.317 | ±0.46 | 7.988 | ±0.83 | 20.545 | ±2.23 | 5.242 | ±0.67 |
| | | *35S::NTT-GR* | 68.734** | ±5.26 | 1.983 | ±0.22 | 5.796* | ±0.66 | 20.330*** | ±1.45 | 35.212* | ±4.03 | 5.414 | ±0.67 |

### trans-Zeatin-types:

| Tissue | Treatment | Genotype | Total tZ-types | SE | tZ | SE | tZR | SE | tZRMP | SE | tZOG | SE | tZROG | SE | tZ7G | SE | tZ9G | SE |
|---|---|---|---|---|---|---|---|---|---|---|---|---|---|---|---|---|---|---|
| Shoot | Mock | **Col-0** | 25.914 | ±1.35 | 0.795 | ±0.09 | 0.927 | ±0.03 | 5.511 | ±0.86 | 3.362 | ±0.15 | 0.156 | ±0.02 | 9.475 | ±0.27 | 5.689 | ±0.24 |
| | | ***35S::NTT-GR*** | 25.343 | ±1.55 | 0.668 | ±0.04 | 0.722 | ±0.12 | 4.291 | ±0.42 | 3.834 | ±0.28 | 0.157 | ±0.00 | 10.025 | ±0.66 | 5.646 | ±0.34 |
| | DEX | **Col-0** | 28.103 | ±1.73 | 2.314 | ±0.35 | 0.650 | ±0.10 | 4.344 | ±0.49 | 4.106 | ±0.48 | 0.201 | ±0.01 | 10.197 | ±0.60 | 6.290 | ±0.31 |
| | | ***35S::NTT-GR*** | 36.355 | ±4.18 | 2.269 | ±0.34 | 1.244 | ±0.20 | 5.664 | ±0.89 | 5.568 | ±0.53 | 0.390* | ±0.06 | 13.586 | ±1.99 | 7.635 | ±1.02 |
| Root | Mock | **Col-0** | 165.121 | ±14.11 | 1.878 | ±0.29 | 5.128 | ±0.79 | 5.870 | ±0.77 | 46.376 | ±6.37 | 0.813 | ±0.11 | 46.894 | ±4.72 | 59.629 | ±2.53 |
| | | ***35S::NTT-GR*** | 155.493 | ±19.32 | 1.217 | ±0.19 | 3.351 | ±0.51 | 5.583 | ±0.50 | 55.818 | ±7.97 | 1.042 | ±0.13 | 39.538 | ±6.04 | 50.341 | ±8.16 |
| | DEX | **Col-0** | 133.035 | ±12.64 | 1.081 | ±0.09 | 3.596 | ±0.37 | 4.891 | ±0.72 | 39.519 | ±5.74 | 0.649 | ±0.09 | 35.755 | ±3.75 | 48.766 | ±6.12 |
| | | ***35S::NTT-GR*** | 306.319** | ±32.25 | 2.169* | ±0.28 | 11.487* | ±2.03 | 14.699* | ±1.73 | 74.744* | ±6.67 | 2.111* | ±0.35 | 75.194* | ±9.34 | 125.916** | ±16.74 |

### cis-Zeatin-types:

| Tissue | Treatment | Genotype | Total cZ-types | SE | cZ | SE | cZR | SE | cZRMP | SE | cZOG | SE | cZROG | SE | cZ7G | SE | cZ9G | SE |
|---|---|---|---|---|---|---|---|---|---|---|---|---|---|---|---|---|---|---|
| Shoot | Mock | **Col-0** | 18.036 | ±1.22 | 0.164 | ±0.02 | 0.288 | ±0.03 | 4.354 | ±0.72 | 0.708 | ±0.10 | 2.548 | ±0.13 | 9.491 | ±0.43 | 0.484 | ±0.02 |
| | | ***35S::NTT-GR*** | 18.856 | ±0.85 | 0.153 | ±0.01 | 0.375 | ±0.05 | 5.286 | ±0.58 | 0.703 | ±0.04 | 2.565 | ±0.15 | 9.290 | ±0.50 | 0.483 | ±0.03 |
| | DEX | **Col-0** | 19.335 | ±0.59 | 0.239 | ±0.04 | 0.463 | ±0.07 | 5.751 | ±0.61 | 0.769 | ±0.06 | 2.552 | ±0.17 | 9.080 | ±0.62 | 0.481 | ±0.03 |
| | | ***35S::NTT-GR*** | 21.092 | ±2.30 | 0.199 | ±0.03 | 0.723* | ±0.04 | 7.339 | ±1.14 | 0.723 | ±0.03 | 3.348 | ±0.47 | 8.287 | ±0.77 | 0.472 | ±0.05 |
| Root | Mock | **Col-0** | 108.291 | ±11.42 | 3.872 | ±0.59 | 9.380 | ±1.00 | 25.353 | ±3.25 | <LOD | <LOD | 2.961 | ±0.46 | 56.354 | ±6.86 | 10.371 | ±0.72 |
| | | ***35S::NTT-GR*** | 93.354 | ±11.36 | 3.244 | ±0.41 | 9.954 | ±1.23 | 27.024 | ±3.96 | <LOD | <LOD | 4.145 | ±0.60 | 41.536 | ±5.99 | 7.450 | ±1.08 |
| | DEX | **Col-0** | 78.105 | ±10.07 | 3.304 | ±0.40 | 6.276 | ±0.89 | 19.637 | ±2.96 | <LOD | <LOD | 2.809 | ±0.40 | 37.993 | ±5.70 | 8.087 | ±1.10 |
| | | ***35S::NTT-GR*** | 105.381 | ±11.42 | 3.947 | ±0.53 | 7.262 | ±0.62 | 25.306 | ±3.37 | <LOD | <LOD | 5.254 | ±0.80 | 51.347 | ±6.77 | 12.264 | ±1.56 |

### Dihydrozeatin-types:

| Tissue | Treatment | Genotype | Total DHZ-types | SE | DHZ | SE | DHZR | SE | DHZRMP | SE | DHZOG | SE | DHZROG | SE | DHZ7G | SE | DHZ9G | SE |
|---|---|---|---|---|---|---|---|---|---|---|---|---|---|---|---|---|---|---|
| Shoot | Mock | **Col-0** | 2.637 | ±0.12 | 0.315 | ±0.04 | 0.122 | ±0.01 | <LOD | <LOD | 0.087 | ±0.00 | <LOD | <LOD | 1.943 | ±0.07 | 0.170 | ±0.02 |
| | | ***35S::NTT-GR*** | 2.463 | ±0.10 | 0.221 | ±0.01 | 0.097 | ±0.01 | <LOD | <LOD | 0.087 | ±0.00 | <LOD | <LOD | 1.903 | ±0.09 | 0.154 | ±0.01 |
| | DEX | **Col-0** | 3.261 | ±0.21 | 0.806 | ±0.12 | 0.117 | ±0.01 | <LOD | <LOD | 0.093 | ±0.01 | <LOD | <LOD | 2.041 | ±0.17 | 0.205 | ±0.01 |
| | | ***35S::NTT-GR*** | 5.353* | ±0.51 | 0.886 | ±0.13 | 0.174 | ±0.02 | <LOD | <LOD | 0.205** | ±0.02 | <LOD | <LOD | 3.853* | ±0.55 | 0.233 | ±0.03 |
| Root | Mock | **Col-0** | 19.846 | ±2.36 | 0.560 | ±0.06 | 0.641 | ±0.10 | <LOD | <LOD | 1.160 | ±0.15 | <LOD | <LOD | 14.230 | ±1.86 | 3.254 | ±0.34 |
| | | ***35S::NTT-GR*** | 19.276 | ±3.66 | 0.380 | ±0.06 | 0.523 | ±0.06 | <LOD | <LOD | 1.414 | ±0.22 | <LOD | <LOD | 12.520 | ±2.72 | 4.438 | ±0.68 |
| | DEX | **Col-0** | 12.488 | ±1.50 | 0.216 | ±0.02 | 0.471 | ±0.06 | <LOD | <LOD | 0.718 | ±0.07 | <LOD | <LOD | 8.569 | ±1.08 | 2.514 | ±0.31 |
| | | ***35S::NTT-GR*** | 35.593** | ±4.91 | 0.697** | ±0.09 | 1.998** | ±0.30 | <LOD | <LOD | 1.947** | ±0.19 | <LOD | <LOD | 24.576** | ±3.52 | 6.375 | ±1.34 |
