## Supplemental Table S1 for "The transcription factor NO TRANSMITTING TRACT / WIP2 regulates cytokinin homeostasis in Arabidopsis"

**Table S1. Primers used in this work**

**Primers for genotyping.**

| **Primer** | **Sequence (5´-3')** |
|---|---|
| 35S Forward | CCTTAATTAAAACCCTCTCCAAATGAAATGAAC |
| 35S Reverse | CCGGTACCATTCTACTCCAAAAATATCAAAGATACAG |
| GFP Forward | ATGCCTGAGGGATACGTGC |
| GFP Reverse | GTGGTCTCTCTTTTCGTTGGG |
| GABI_363C02 (*ckx7)* Rp | GAACATCAGAATCTTCCACCG |
| GABI_363C02 (*ckx7)* Lp | GAACTGTGTGAAGCCTCTTGC |

**Primers for RT-PCR.**

| **Primer** | **Sequence (5´-3')** |
|---|---|
| Actin2 F | AATCACAGCACTTGCACC |
| Actin2 R | AATCACAGCACTTGCACC |
| CKX7 F | ACTCCGATTCCAACTCAACC |
| CKX7 R | CGATTTCTTCTGACCGTTTG |
| IPT5 F | CGATGACGAAAGAAGGGAAG |
| IPT5 R | CTCCAAGACAGCGACCAATC |
| AHP6 F | TAACGTCTGCGTTGCCTTT |
| AHP6 R | CCTCCAGTCCTCTCAAGCAC |

**Primers for cloning the Y1H promoters and fragments.**

| **Primer** | **Sequence (5´-3')** |
|---|---|
| CKX7 F | CGTGGTCAAAAAGAGAAAACTTG |
| CKX7 R | TTGATAGAATGGCCGTTTCC |
| IPT5 F | CCGACTCCGATCAAACATGC |
| IPT5 R | TCGAGCTCTGGAACTCCAAT |
| AHP6 F | CATCTCAATGACTCATCATATCGAATGT |
| AHP6 R | CCACAACGGCACACCCGT |
| PUP10 F | CCTCCTCGTGGTCAACACTGA |
| PUP10 R | CATGGTGGATGTGGAATCAC |
| KAN1 F | TTAGGTTCTTTAGGGTTCACTTGTTT |
| KAN1 R | CATAATTAAAGAAACCTTTCTCTTGC |
| IPT5 A F | CAGTAACATTCCTCCAGCC |
| IPT5 A R | GCTCTCAACGAGCATATG |
| IPT5 B F | TGTCTCGGAGCGAGTTGATA |
| IPT B R | GAAGGCGCCAATATTCTCCT |
| AHP6 A F | GTGAAACAATTTACG |
| AHP6 A R | TTAATTTCTCACTGTTGAAATTAATCTC |
