## Supplementary Figures S1-S7 for "The transcription factor NO TRANSMITTING TRACT / WIP2 regulates cytokinin homeostasis in Arabidopsis"

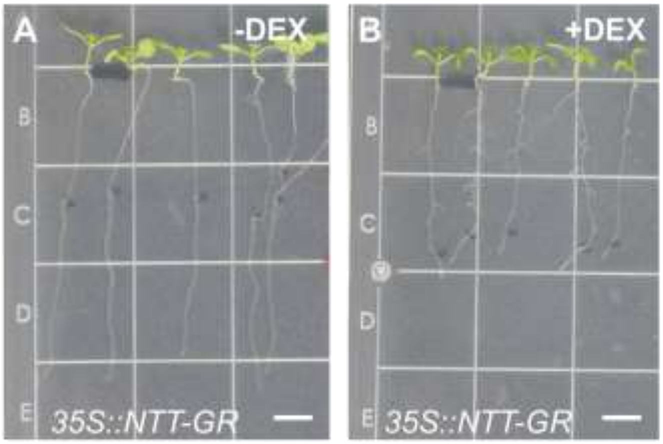


**Figure S1. Root phenotype of induced *35S::NTT-GR* seedlings.** (A) Mock-treated 4 dag seedlings, shown 3 days after induction. The root length at the time of induction is marked by a black dot on the plate. (B) DEX-treated seedlings (same time after induction as in A) show clearly diminished growth.

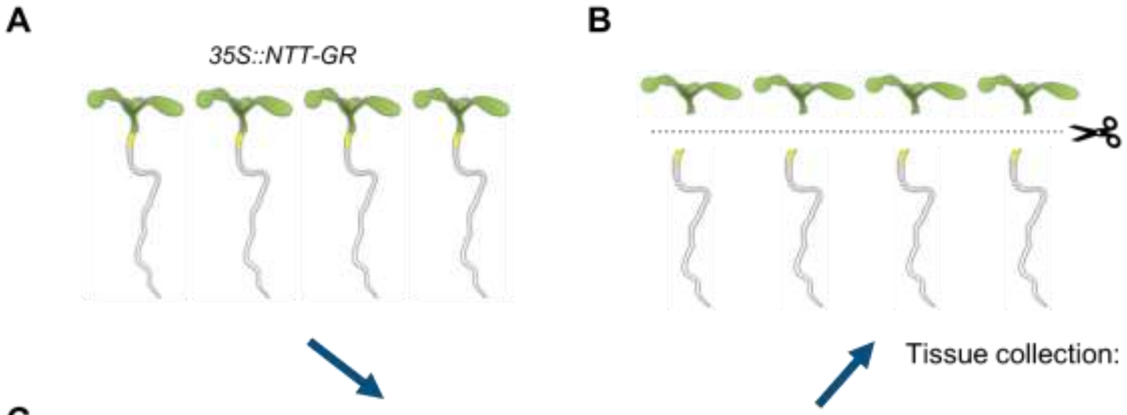


C

| Treatment | Ethanol | DEX (100 μM) | CHX (30 μM) |
|---|---|---|---|
| Induction 30 m | + | + | + |
| Induction 8 h | + | + | |
| Control 30 m | + | | + |
| Control 8 h | + | | |

**Figure S2. Global expression experiment workflow.** (A) *35S::NTT-GR* 4 dag seedlings were treated according to the table in C. (B) After the designated time points (30 min or 8 h), aerial and root tissues were collected separately. (C) Treatment table.

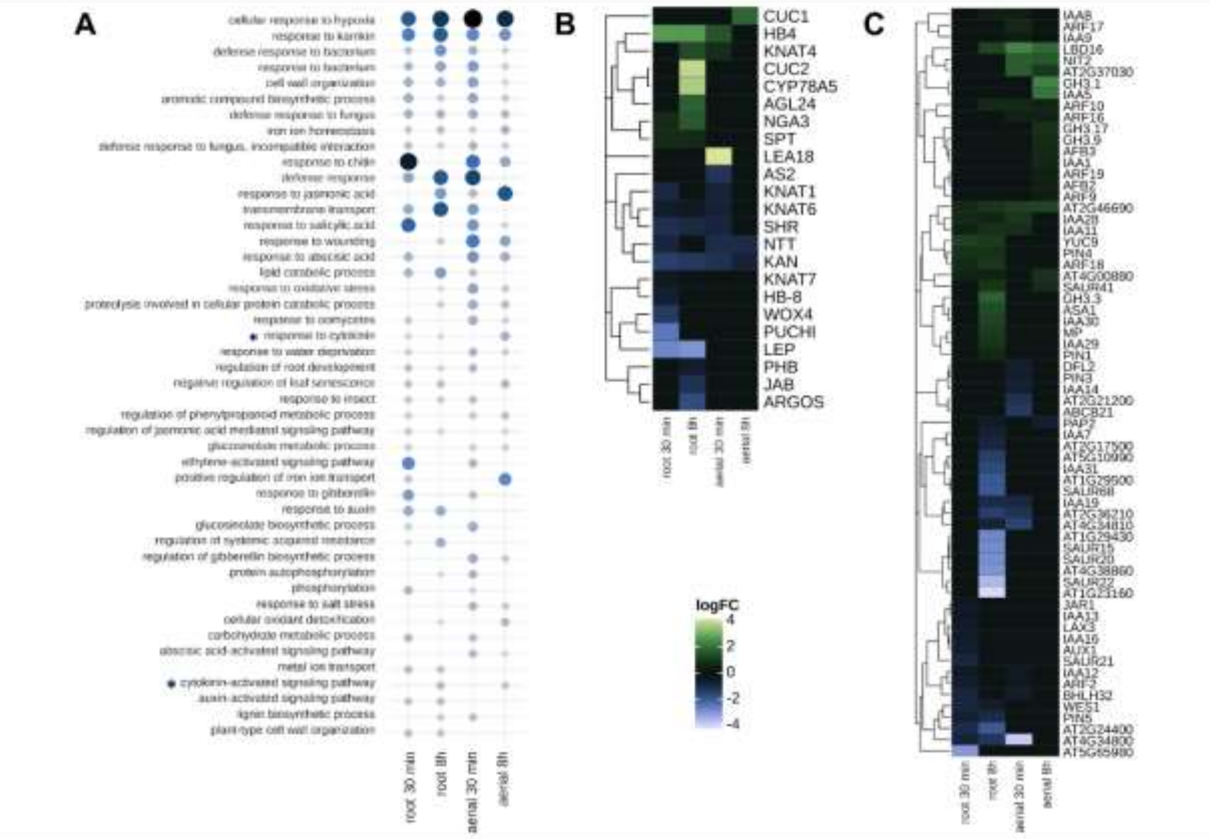


**Figure S3. Bubble plot of GO enriched categories and Heatmap of developmental and auxin-related differentially expressed genes.** (A) Bubble plot showing the most enriched Gene Ontology (GO) categories. Bubble size and color intensity represent the FDR (False Discovery Rate), with larger, darker bubbles indicating lower FDR values. (B) Heatmap of genes associated with plant development regulators (C) Heatmap of auxin-related genes showing upregulated and downregulated genes across conditions. The color scale represents $\log_2$ fold-change.

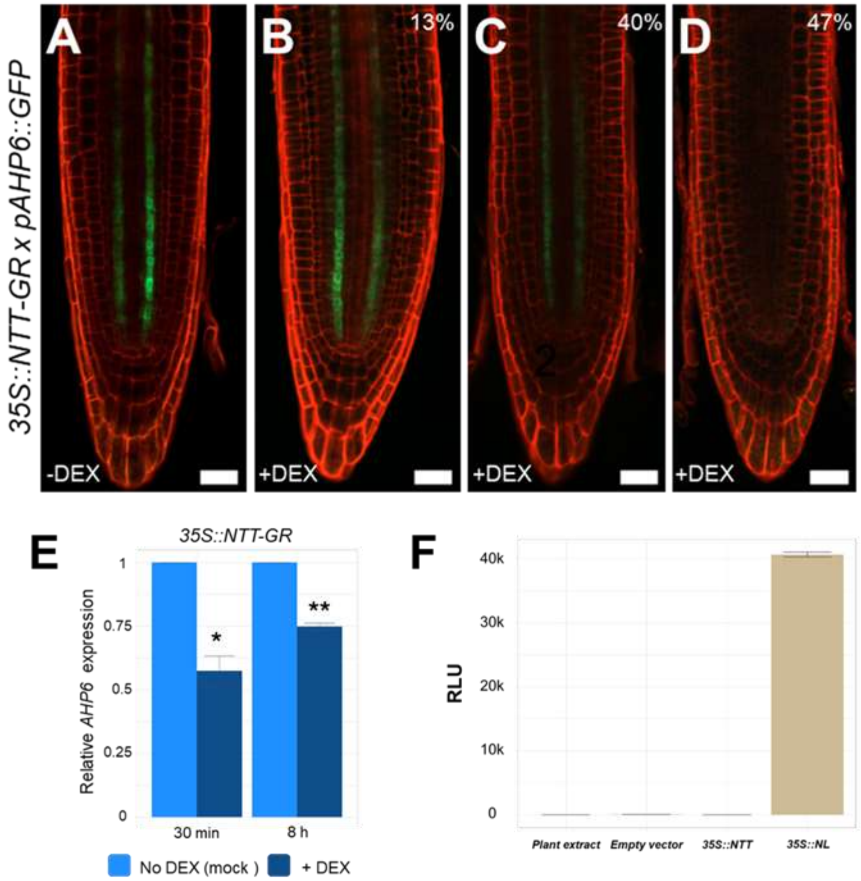


**Figure S4. Change in expression of *AHP6* after NTT induction.** 4 dag *35S::NTT-GR* x *pAHP6::GFP* seedlings were treated and analyzed by confocal microscopy 48 h after induction (2 dai). (A) Mock treated root, with no change in expression compared to the *pAHP6::GFP* line in the wild type background. (B-D) DEX treated seedlings. (B) 13% of roots displayed normal expression, comparable to mock-treated roots. (C) 40% showed a decreased GFP signal, compared to mock treated roots. (D) In 47% of roots, no GFP signal was detected, in contrast to the strong signal observed in ~80% of mock-treated seedlings (due to segregation). Scale bars: 25 µm. Red corresponds to propidium iodide staining, and green to GFP signal. (E) *AHP6* relative expression at 30 min and 8 h after NTT induction, in induced (+DEX, dark blue) and mock treated (-DEX, light blue) root tissue. The y-axis represents fold change. Three biological replicates were used for each measurement. Values are given as mean ± SE, and asterisks indicate significant differences between conditions (* $p<0.05$, ** $p < 0.01$, Student's t-test). (F) NanoLuc assay controls. The x-axis indicates the different controls used (plant extract, empty vector, effector only, and *35S::NL*). The y-axis: represent Relative Luminescence Units (RLU).

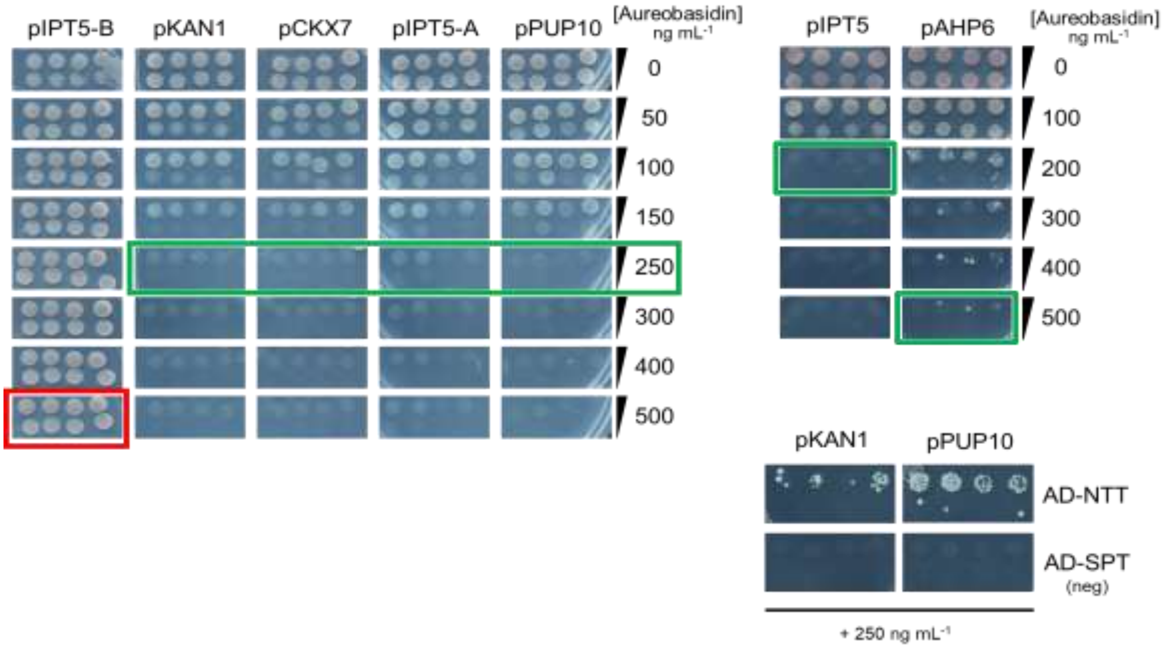


**Figure S5. Y1H Autoactivation assays, and *pKAN* and *pPUP10* experiments.** Autoactivation assays were performed using different concentrations of Aureobasidin, to choose the right concentration for the Y1H experiments. One of the *IPT5* fragments (*IPT5-B*, corresponding to the internal sequence) consistently showed autoactivation and was therefore unsuitable for Y1H experiments. The promoters of *KAN* and *PUP10* were also tested. The results indicate that NTT binds to them, whereas *SPATULA (SPT),* used as a negative control, does not.

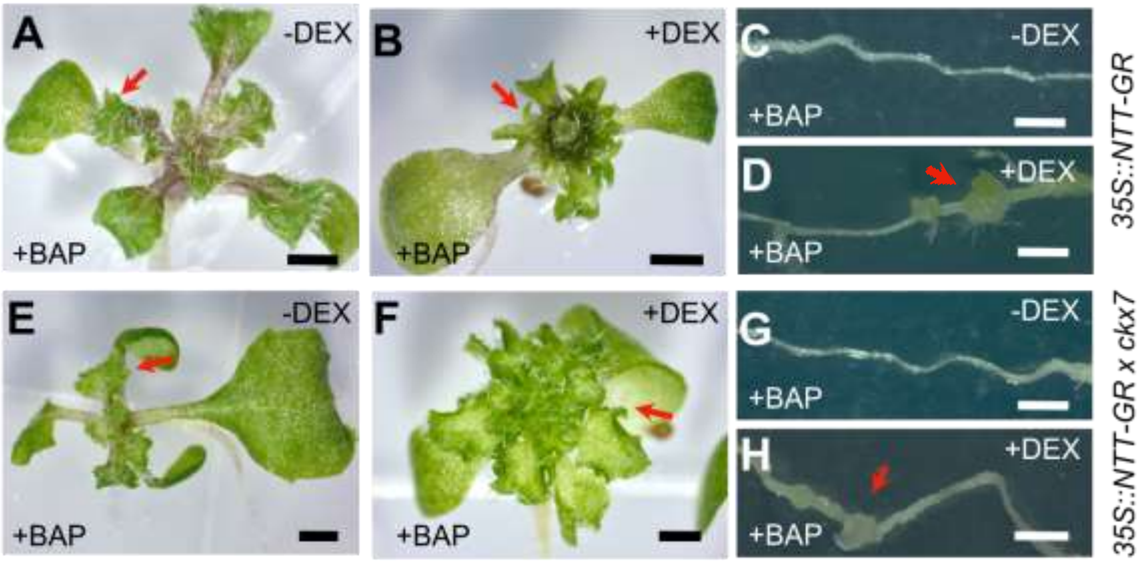


**Fig. S6. NTT induction phenotype in *ckx7* seedlings, treated with cytokinin.** Shoots of *35S::NTT-GR* (A,B) and *35S::NTT-GR* x *ckx7* (E,F) seedlings in BAP supplemented medium, either mock treated (A,E), or induced with DEX (B,F); Roots of *35S::NTT-GR* (C,D) and *35S::NTT-GR* x *ckx7* (G,H) seedlings in BAP supplemented medium, either mock treated (C,G) or induced with DEX (D,H). Red arrows indicate irregularities in the leaf margins or callus formation in roots. Scale bars represent 1 mm in all cases.

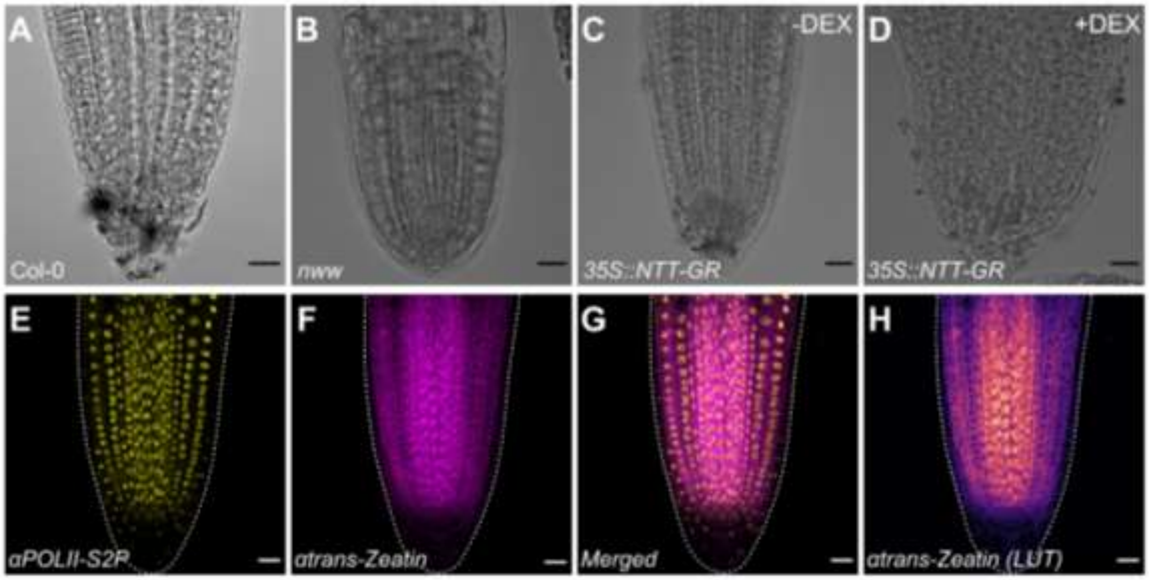


**Figure S7. Root phenotypes and immunolocalization technical control.** (A-D) Representative brightfield images showing root tip phenotypes of 1 dag seedlings in (A) Col-0, (B) *nww* and in (C, D) *35S::NTT-GR*, either mock-treated (C) or induced with DEX (D). (E) Localization of Pol II S2p, used as a technical control. (F) Representative image of the localization of trans-Zeatin-riboside, without LUT transformation. (G) Merged image of Pol II S2p and trans-Zeatin-riboside localization. (H) LUT transformed image of trans-Zeatin-riboside localization, enhancing signal visualization. Scale bars: 20 µm.
